# High-resolution cryo-EM analysis of the therapeutic *Pseudomonas* phage Pa223

**DOI:** 10.1101/2025.06.11.659172

**Authors:** Chun-Feng David Hou, Nathan Bellis, Ravi K Lokareddy, Steven Branston, Johnny Reid, Renae Geier, Angela Soriaga, Lucy Sim, Pierre Kyme, Deborah Birx, Sebastien Lemire, Gino Cingolani

**Affiliations:** Department of Chemistry and Chemical Biology, Rutgers, the State University of New Jersey, Piscataway, NJ, 08854, USA; Department of Biochemistry and Molecular Genetics, University of Alabama at Birmingham, 1825 University Blvd, Birmingham, AL 35294, USA; Armata Pharmaceuticals Inc., 5005 McConnell Ave, Los Angeles, CA 90066, USA

**Author notes:** **Corresponding Author:** Gino Cingolani, Ph.D., Department of Biochemistry and Molecular Genetics, Univ of Alabama at Birmingham, 1825 University Blvd, Birmingham, AL 35294, USA.

**Keywords:** *Pseudomonas* phages, Bryunoghevirus, PaP3-like phage, cryo-EM, ejection proteins, genome ejection, cystic fibrosis, phage therapy

## Abstract

Cryogenic electron microscopy (cryo-EM) analysis of bacteriophages is a valuable method for deciphering virus composition and conformational plasticity. In this study, we present a high-resolution structural atlas of the *Pseudomonas* virus Pa223, a phage from the *Bruynoghevirus* genus that has recently been used in clinical cocktails for treating cystic fibrosis and non-cystic fibrosis bronchiectasis, as well as for compassionate care. By combining bioinformatics, proteomics, cryo-EM single particle analysis, and localized reconstruction, we annotated and built atomic models for eight structural polypeptide chains that form the icosahedral capsid and noncontractile tail. We discovered that the Pa223 capsid is decorated by a spike protein that features a unique triple-β helix fold with no structural homologs in the database. The Pa223 tail features six trimeric tail fibers extending upwards, similar to, but shorter than, those found in phage T7. Unlike T7, the Pa223 tail is extended by two head-to-tail adaptors and sealed by a trimeric tail needle, similar to P22-like phages. We identified a protein bound around the outer perimeter of the portal protein, positioned similarly to the ejection protein gp72, which was identified in the *Pseudomonas* phage DEV, a *Litunavirus* phage and member of the reclassified *Schitoviridae* family. This structural hint led us to identify the Pa223 ejection proteins gp53, gp54, and gp56, which bioinformatically resemble those of T7-like phages more closely than *Schitoviridae*. Thus, phage Pa223 contains diverse structural elements found in P22-like, T7-like, and *Litunavirus* phages, providing a framework for understanding the diversification and evolution of ejection proteins in *Bruynogheviruses*.

**Highlights:**

- The high-resolution structure of *Bruynoghevirus* Pa223 reveals hybrid structural features that are shared among P22-like, T7-like, and *Litunavirus* phages.
- The Pa223 capsid is decorated with a trimeric spike asymmetrically bound at the icosahedral 3-fold axes.
- The Pa223 tail features two quasi-equivalent conformations of the head-to-tail adaptor protein arranged into two coaxial rings.
- Identification of the ejection protein gp54 through structural similarity to gp72 from the *Litunavirus* DEV.
- Bioinformatic mapping of the Pa223 ejection proteins gp53 and gp56 validated through mass spectrometry analysis of infectious virions.

## INTRODUCTION

The Gram-negative pathogen *Pseudomonas aeruginosa* is a major cause of morbidity and mortality worldwide [1]. Infections caused by *P. aeruginosa* are significant in cystic fibrosis (CF) patients, who experience elevated mucus levels in the lungs and impaired mucociliary clearance of microbial pathogens, including *P. aeruginosa* [2, 3]. Recurring *P. aeruginosa* infections and persistent inflammation are particularly concerning in adult CF patients [4], whose bacterial colonies develop specific mutations and adaptive responses to antibiotic exposure, leading to the selection and spread of multidrug-resistant (MDR) strains [5], which are often linked to antibiotic-impermeable biofilms. Phage therapy aimed at *P. aeruginosa* offers a promising therapeutic approach for tackling CF-related infections [6, 7]. Phages are advantageous because they are effective against both antibiotic-sensitive and antibiotic-resistant strains of *P. aeruginosa*, generating renewed interest in discovering, characterizing, and engineering *Pseudomonas* phages for therapeutic applications.

Pa223 is a *P. aeruginosa* lytic bacteriophage of the *Bruynoghevirus* genus [8], characterized by a short, noncontractile tail similar to the coliphage T7 [9, 10]. Phages from the *Bruynoghevirus* genus have been utilized in experimental phage therapy cocktails due to their broad host range and high lytic activity against both planktonic and biofilm-associated bacteria [8, 11, 12]. Bioinformatics analysis indicates that these small phages do not encode any known bacterial toxins or virulence factors that could pose a risk to humans. Consequently, Pa223 is included in a therapeutic cocktail of five phages used by Armata Pharmaceuticals to treat *P. aeruginosa* infections in CF patients and those with non-CF bronchiectasis as part of clinical trials SWARM-P.a. (NCT04596319) and Tailwind (NCT05616221), respectively.

Pa223 possesses a double-stranded DNA (dsDNA) genome of approximately 45.7 kbp that encodes 71 open reading frames (ORFs). At the genome level, Pa223 is 72.3% identical to *Pseudomonas* phage 3 (PaP3) [13] and 83.6% identical to the intron-containing phage LUZ24 [14], while PaP3 and LUZ24 are 70.2% identical. All three phages, Pa223, PaP3, and LUZ24, are *Bruynogheviruses*. Pa223 encodes a large terminase subunit (TerL) that contains the classical domain signature, which includes an N-terminal ATPase and a C-terminal nuclease domain [15]. Meanwhile, Pa223 TerS is 89% identical (>95% similar over 152 residues) to the TerS of PaP3, which is nonameric; it binds the *cos* packaging initiation site and stimulates the ATPase activity of TerL [15, 16].

Many podophages have been characterized at near-atomic resolution in recent years, leveraging cryo-EM and bioinformatic and proteomic analysis. Among them are the P22-like phages Sf6 [17], CUS-3 [18], and P22 [19]; T7-like phages [9, 10], HRP29 [20], G4 [21], Pam1 [22] and P-SCSP1u [23]; the marine phage Carin-1 [24]; phi29 [25] and *Staphylococcus* phages [26–28]. Despite this progress, annotating *Pseudomonas* phage proteins remains challenging and inherently inaccurate without structural information due to extensive gene and cassette swapping among phages. In this study, we describe the atomic structure of the *Pseudomonas* phage Pa223, solved using cryo-EM Single Particle Analysis (SPA) and localized reconstruction, validated by liquid chromatography mass spectrometry (LC-MS/MS) analysis of the purified Pa223 virion and bioinformatic analysis of its genome. The high-resolution structural atlas presented here highlights both the structural simplicity and mosaic composition of this small phage, which incorporates features observed in several other podophages.

## RESULTS

### Atomic structure of *Pseudomonas* phage Pa223

We used a multi-pronged approach to obtain a high-resolution reconstruction of *Pseudomonas* phage Pa223 (Fig. S1A). First, we generated an asymmetric reconstruction (C1) of the entire virion from 15,698 particles at 4.0 Å resolution, which aligned from C5-C6 symmetry (Fig. S2). This reconstruction, generated without imposing symmetry, was sufficient to discern all symmetric and asymmetric components of the Pa223 virion (Fig. 1A, B). We then calculated four localized reconstructions (Fig. S1B) [29] of the phage capsid (icosahedral 5-fold), capsid spikes (icosahedral 3-fold), protal protein, and tail apparatus that was subjected to different rotational symmetry averaging, yielded density maps for all tail factors between 2.6 and 3.1 Å resoltuion (Table 1, Fig. S2). The tail needle, the least visible feature in the reconstruction, was deciphered using a focus reconstruction of the tail tip that yielded a complete map to 4.1 Å resolution (Table 1, Fig. S2). In all cases, the reconstructions had excellent side-chain density, allowing us to annotate and *de novo* build eight Pa223 ORFs (Fig. 1A, B), which were refined to a map-to-model correlation coefficient (CC) between 0.7 and 0.91 (Table 1). Pa223’s structural atlas presented in this paper (Fig.1A, B) comprises seven structural proteins, three in the capsid: gp65 (coat protein), gp52 (capsid spike), and gp68 (portal protein) and four in the short tail apparatus: gp63 (head-to-tail adaptor), gp60 (tail fiber), gp59 (tail hub), gp58 (tail needle). In addition, we identified the ejection protein gp54 and mapped gp53 and gp56 as putative ejection proteins invisible by cryo-EM but present inside the capsid. Liquid chromatography-mass spectrometry (LC-MS) analysis of virions used for cryo-EM analysis confirmed the existence of all proteins modeled in the cryo-EM density and mapped bioinformatically (gp53 and gp56) and identified an additional ∼11 kDa protein, gp31, not seen in the reconstruction (Table 2).

**Fig. 1.**
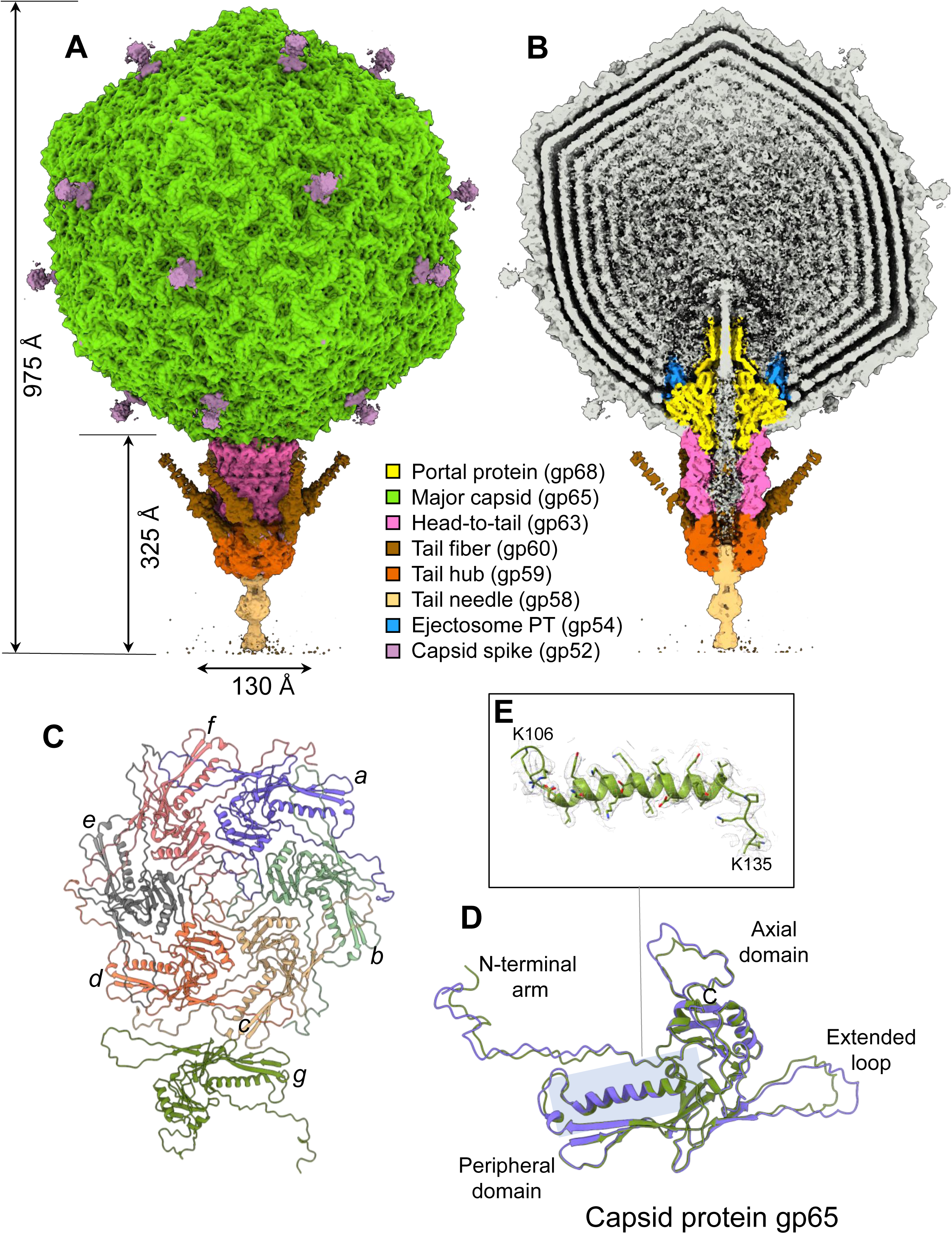
Cryo-EM reconstruction of *Pseudomonas* phage Pa223. Side (A) and cut-out (B) view of Pa223 asymmetric reconstruction at 4.0 Å resolution. (C) Pa223 capsid icosahedral unit refined at 2.6 Å resolution. (D) Overlay of capsid protein conformers found in the penton (green) and hexon (blue). (E) A representative 2.6 Å density of Pa223 capsid protein contoured at 5σ (light gray) overlaid to the refined model for residues 106-135 (green).

**Table 1.** Cryo-EM models refinement statistics.

| Data Collection |  | Pseudomonas phage Pa223 |  |  |  |  |  |  |  |
| --- | --- | --- | --- | --- | --- | --- | --- | --- | --- |
| Facility |  | Stanford-SLAC CryoEM Center (S<sup>2</sup>C<sup>2</sup>) |  |  |  |  |  |  |  |
| Microscope/Camera |  | Titan Krios/Falcon 4 |  |  |  |  |  |  |  |
| Magnification |  | 150,000X |  |  |  |  |  |  |  |
| Voltage (kV) |  | 300 |  |  |  |  |  |  |  |
| Electron exposure (e<sup>-</sup>/Å<sup>2</sup>) |  | 50 |  |  |  |  |  |  |  |
| Defocus range (um) |  | -0.8 to -1.6 (step 0.2) |  |  |  |  |  |  |  |
| Pixel size (Å) |  | 0.946 |  |  |  |  |  |  |  |
| Tot movies (frames/movie) |  | 13,327 (40) |  |  |  |  |  |  |  |
| Processing and Refinement |  |  |  |  |  |  |  |  |  |
| PDB entry code |  | 9NY2 | 9NWI / 9NXK |  |  | 9NWM / 9NXO |  | 9NXP | 9NY6 |
| EMDB entry code |  | EMD-49916 / EMD-49926 | EMD-49882 / EMD-49902 |  |  | EMD-49887 / EMD-49909 |  | EMD-49910 | EMD-49925 |
| Particles per reconstruction |  | 166,546/15,698 | 46,151/80.543 |  |  | 46,120 |  | 14,155 | 382,465 |
| Map Symmetry |  | C5/C1(bin2) | C6/C1(focused) |  |  | C12/C12 (PT-focused) |  | C3 | C1 |
| Model name |  | Capsid Protein/ Full virion | HT-adaptor+Tail Hub/ Tail Fiber |  |  | Portal/ Ejection Protein |  | Tail Needle | Capsid spike |
| Map resolution (Å) |  | 2.6/4.0 | 2.8/3.1 |  |  | 2.6/3.0 |  | 4.1 | 3.0 |
| FSC threshold |  | 0.143 |  |  |  |  |  |  |  |
| Initial model used |  | de novo | denovo/AlphaFold |  |  | PDB:7SYA/de novo |  | de novo | de novo |
| Model to Map Correlation Coefficient (CC) |  | 0.91 | 0.91/0.86 |  |  | 0.88/0.78 |  | 0.70 | 0.82 |
| Model composition | Number of chains | 7 | 24 | 6 | 3 | 12 | 12 | 3 | 3 |
|  | Non-H atoms | 17,199 | 40,704 | 24,390 | 4,653 | 60,720 | 13,296 | 5,304 | 6,705 |
|  | Residues | 2,219 | 4,968 | 3,072 | 642 | 7,536 | 1,752 | 684 | 885 |
| RMS deviations | Bond lengths (Å) | 0.003 (0) | 0.004 (0) | 0.005 (0) | 0.003 (0) | 0.002 (0) | 0.003 (0) | 0.003 (0) | 0.003 (0) |
|  | Bond angles (°) | 0.529 (0) | 0.504 (1) | 0.582 (0) | 0.570 (0) | 0.493 (0) | 0.675 (0) | 0.727 (0) | 0.608 (1) |
| Validation | MolProbity score | 1.7 | 1.5 | 1.8 | 2.0 | 1.8 | 2.1 | 2.2 | 2.0 |
|  | ClashScore | 5.3 | 3.8 | 6.5 | 14.8 | 5.6 | 9.3 | 16.5 | 13.7 |
|  | Rotamer outliers % | 1.47 | 0.19 | 0.11 | 0.00 | 1.62 | 0.75 | 0.00 | 0.14 |
| Ramachandran plot | Favored % | 96.01 | 95.57 | 92.22 | 94.81 | 95.24 | 93.29 | 92.92 | 94.54 |
|  | Allowed % | 3.58 | 3.94 | 7.39 | 5.19 | 4.29 | 6.71 | 7.08 | 5.35 |
|  | Outliers % | 0.41 | 0.49 | 0.39 | 0.0 | 0.48 | 0.00 | 0.00 | 0.11 |

**Table 2.** Identification of Pa223 Major Structural Proteins by LC-MS/MS.

| **Protein** | **Mascot**<br>**Score<sup>a</sup>** | **MW (Da)** | **Annotation** |
| --- | --- | --- | --- |
| gp065 | 11,898 | 34,990 | Major Capsid Protein |
| gp052 | 5,400 | 31,676 | Capsid Spike |
| gp054 | 4,988 | 53,516 | putative Ejection Protein |
| gp053 | 2,220 | 112,061 | putative Ejection Protein |
| gp068 | 2,007 | 81,027 | Portal Protein |
| gp059 | 1,130 | 57,630 | Tail Hub |
| gp056 | 1,065 | 32,038 | putative Ejection Protein |
| gp063 | 539 | 24,071 | Head-to-Tail Adaptor |
| gp060 | 328 | 22,125 | Tail Fiber |
| gp031 | 187 | 10,934 | unknown |
| gp058 | 56 | 25,133 | Tail Needle |

a. Statistical measurement of how well the observed MS data matches the identified protein sequence; scores greater than 50 represent correct identifications and can reflect relative protein abundance within a sample.

### Architecture of *Pseudomonas* phage Pa223 T = 7 capsid

The Pa223 virion has a maximum length of 975 Å, of which the icosahedral capsid accounts for ∼650 Å, while the short noncontractile tail extends the unique vertex by ∼325 Å. The asymmetric reconstruction of the Pa223 capsid at 4.0 Å resolution (Fig. 1A, B) and a 2.6 Å focused reconstruction of the capsid five-fold vertex revealed two components of the Pa223 head: the major capsid protein gp65 (res. 1-317) and the capsid spike gp52 (res. 2-252). The capsid is built by 415 copies of gp65 that form a lattice with a triangulation number T = 7 (Fig. 1C). Gp65 adopts a canonical HK97 fold (Fig. 1D) that comprises an E-loop (residues 52-78), Peripheral domain (P-domain; residues 94-108; 275-288), N-arm (residues 1-33), backbone helix (residues 108-129), and Axial domain (A-domain; residues 131-268, 303-317) [30]. Pa223 capsid protein conforms to the *quasi*-equivalence theory of viral capsids [31], adopting two *quasi*-equivalent conformations in the phage T = 7 capsid: one in the hexon and another in the penton (Fig. 1C, D). The RMSD between these conformers is 0.7 Å, with 246/317 C-alphas aligned. The final model of the icosahedral asymmetric unit was refined to a CC = 0.91 at 2.6 Å resolution (Table 1).

The second protein identified in Pa223 capsid reconstruction is the capsid spike gp52, visible in the asymmetric reconstruction at the three-fold icosahedral symmetry axes as a low-resolution blob (Fig. 1A, B). To identify the gene encoding this protein and build an accurate atomic model, we calculated a localized reconstruction focusing on the capsid 3-fold vertex. We used a cylindrical mask covering one vertex, and after symmetry expansion, we found a class from one of the three orientations representing 24% of the particles with a spike-like protein extending ∼100 Å from the capsid surface. This asymmetric reconstruction yields a 3.0 Å resolution map, allowing us to build a complete atomic model of gp52 (res. 2-295) that was real-space refined to CC = 0.82 (Fig. 2A, B, Table 1). Pa223 capsid spike leans 60° forward relative to the capsid floor. The protein is a homo-trimer comprising three parts: an N-terminal attachment (NTA) domain (res. 2-41), a middle domain comprising a triple β-helix (res. 42-200), and a C-terminal knob (CTK, res. 201-293) (Fig. 2B). A superimposition of gp52 chains (Fig. 2C) reveals a global RMSD of 4.5 Å, with residues 42-293 virtually identical, whereas the N-terminal res. 2-41 are displaced as much as 20 Å from each other.

**Figure 2.**
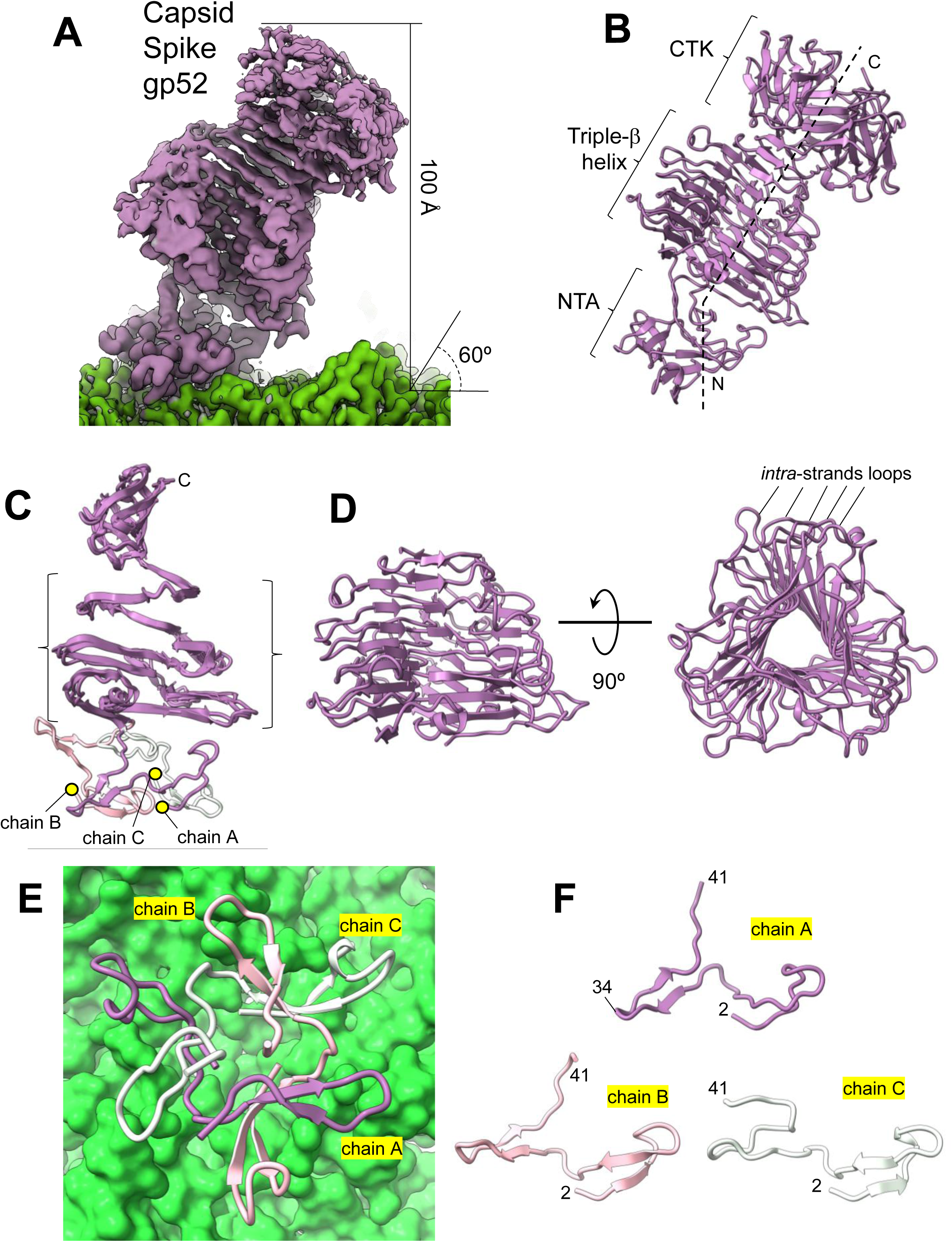
Reconstruction of Pa223 capsid spike. (A) Focused reconstruction of the capsid spike 52 (purple) overlaid to the capsid density (green) at 3.0 Å resolution. (B) Ribbon diagram of the capsid spike gp52 (purple), with its three domains labeled as N-terminal attachment (NTA), triple β-helix, and C-terminal knob (CTK). (C) Superimposition of gp52 protomers (chains A, B, and C) reveals asymmetry of N-terminal residues 1-41 (colored purple, pink, and white, respectively). (D) Ribbon diagram of gp52 triple-β helix displayed in top and side views. (E) Interaction of the capsid spike N-terminal residues 2-41 (colored purple, pink, and white) with the capsid proteins (shown as a green solvent surface). (F) Ribbon diagrams of the capsid spike residues 2-41 from chains A (purple), B (pink), and C (white) are displayed in the same orientation. Residues 2-34 of chain A make the most contact with the capsid floor.

Interestingly, the Pa223 capsid spike lacks structural homology to any decorating protein visualized thus far [32]. Except for the C-terminal knob, which adopts a canonical TNF fold, the gp52 triple β-helix lacks structural similarity to known proteins in the RCSB database, according to Foldseek [33]. It superficially resembles the prism-like structure of phages P22 [19] and Sf6 [34] tailspikes but is more globular due to extended intra-strands loops that significantly expand the helical diameter (Fig. 2D). However, the similarity between the P22-like tailspike and the Pa223 capsid spike is purely morphological and is not backed by statistically significant sequence conservation.

To decipher how the capsid spike is attached to the icosahedral 3-fold axis, we scrutinized the gp52:capsid interaction (Fig. 2E). Unexpectedly, we found that each trimeric spike sits at the capsid 3-fold in a very asymmetric way, making close (< 4 Å) contacts with six capsid proteins. The spike NTA domain (res. 2-41) is solely responsible for anchoring the spike to the capsid, stabilized by 17 hydrogen bonds and six salt bridges. This predominantly polar binding interface, cemented by electrostatic interactions, may contribute to capsid stability. Interestingly, the spike:capsid association is highly asymmetric, with spike chains making a different number of bonds (Fig. 2F). Specifically, chain A makes the most bonds with the capsid surface, namely, eight hydrogen bonds and four salt bridges (through residues E6, D20, R15 and R27) interacting with three capsid proteins. Chain B contacts only two capsid proteins, making just three hydrogen bonds. Finally, chain C makes five hydrogen bonds and two salt bridges (through residues D6 and R15), interacting with just two capsid proteins. Overall, the NTA domain cements the Pa223 spike at the icosahedral 3-fold axes, projecting the spike body outward, which leans 60° relative to the capsid floor.

### Symmetry mismatches in phage Pa223 tail

We generated two high-resolution symmetric reconstructions of the Pa223 tail (Fig. 3A), imposing C12 and C6 symmetry. We identified three symmetry mismatches besides the 12:5 of the portal:capsid [35]. The C12 map was used to identify and build *de novo* models of the portal gp68 and head-to-tail adaptors gp63 and to identify a new factor, gp54. The C6 map gave us atomistic information about the tail hub (gp59) and tail fiber (gp60). Finally, a localized reconstruction helped identify and build the tail needle, gp58, which is 3-fold symmetric.

**Figure 3.**
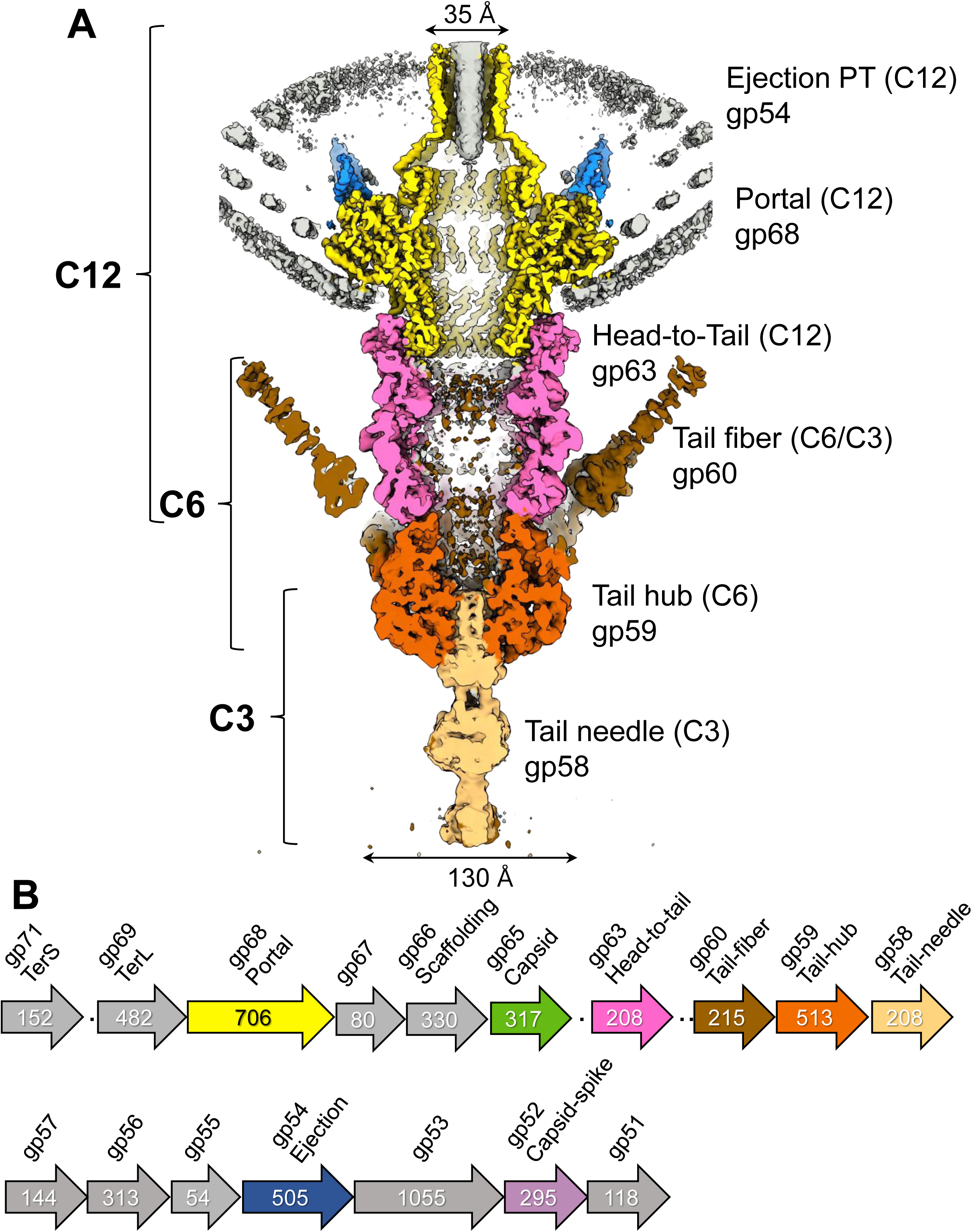
Symmetry mismatches in the Pa223 tail. (A) A magnified, cut-out view of the Pa223 asymmetric reconstruction contoured at 4, colored as shown in Figure 1. The left brackets indicate different degrees of rotational symmetry in the tail components. (B) Schematic diagram of the Pa223 ORFs, color-coded as in panel (A). The number inside each ORF arrow box indicates the predicted number of amino acids.

Pa223 portal protein was built in a 2.6 Å map, and the final model was refined to CC = 0.88 (Fig. 4A, Table 1). The map lacks density for the C-terminal residues 640-706. LC-MS identified these residues that are invisible in the reconstruction, likely due to the symmetry mismatch between the portal and capsid [34] or structural flexibility that prevents alignment. The Pa223 portal is most similar to the PaP3 portal protein, which shares an 84% sequence identity, and was recently solved from recombinantly expressed protein [36]. It presents a classical portal protein fold, including a barrel, wing, stem, and barrel domains (Fig. 4B) [37]. In our reconstruction, the C-terminal barrel forms eight hydrogen bonds between parallel α-helices but does not contain salt bridges. PISA [38] does not predict that the barrel:barrel interface is stable on its own (ΔiG P-value = 0.83), suggesting that the barrel is stabilized by the remaining portal protein structure and the surrounding DNA [39]. Finally, the C12 map revealed strong density features around the portal perimeter (Fig. 4C), which is also visible in the asymmetric reconstruction of the whole virion (Fig. 1B). To improve this density, we calculated a focused reconstruction that allowed us to identify the relative gene product as gp54 (Fig. 3B). We modeled residues 174-319 of this 505 amino acid gene product of unknown function, located downstream of the main structural proteins. The gp54 model was real-space refined against the C12 density to a CC = 0.78 at 3.0 Å resolution (Table 1).

**Figure 4.**
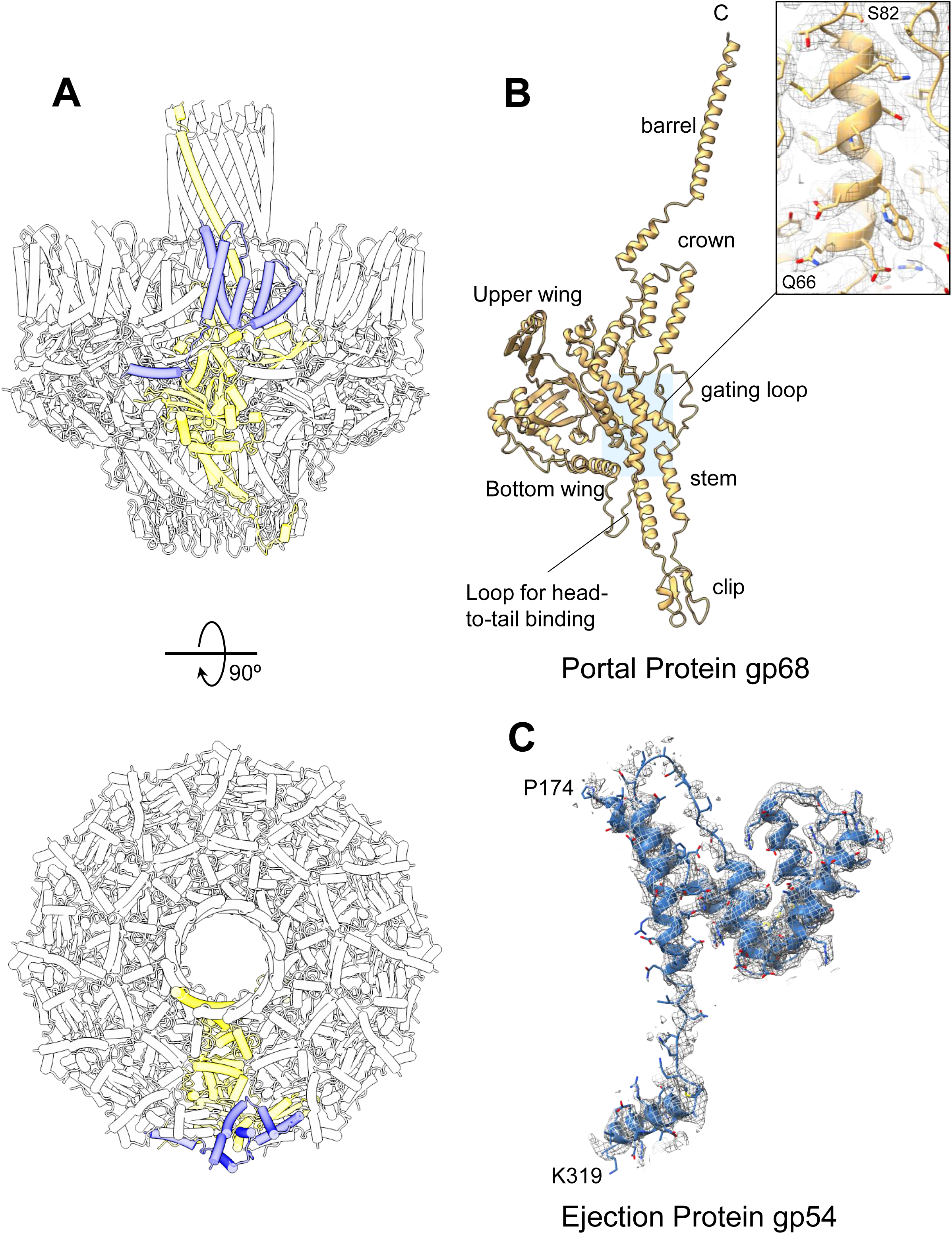
Identification of the Pa223 portal and the putative ejection protein gp54. (A) Side view (top panel) and top view (bottom panel) of the Pa223 portal protein gp68 and gp54, with only one protomer of gp68 and gp54 colored yellow and blue, respectively. All other subunits are shown in light gray; α-helices are represented as cylinders. (B) A magnified view of the portal protein protomer (light orange) includes a zoomed-in view of the 2.6 Å cryo-EM density encompassing residues Q66-S82. (C) A focused reconstruction of gp54 at 3.0 Å resolution is overlaid with the final refined model (blue) from residues P174 to K319.

### Pa223 short-tail composition

Assembled under the portal, we identified two coaxial dodecameric rings formed by the head-to-tail (HT) adaptors, which we named gp63-a and gp63-b (Fig. 5A). We modeled residues 1-208 of either HT-adapt or ring, which were refined to a CC = 0.92 at 2.8 Å resolution. The two HT-adaptor protomers are identical except for the last C-terminal 14 residues 195-208 (Fig. 5B), which adopt a random coil conformation in the conformer gp63-a, bonding the portal protein versus a short 3/10 helix in gp63-b at the interface between the two HT-rings. Secondary structure superimposition reveals an RMSD of 0.6 Å for 194/208 C-alphas, underscoring the *quasi*-equivalence of these two adaptors. Together, the two HT-adaptors extend the portal lumen by 120 Å and have an internal lumen of 50 Å. Pa223 HT-adaptor shares the most significant structural similarity with phage SU10 HT-adaptor, which is 19.7% sequence identical [40] and is significantly different than the HT-adaptor found in P22-like that has a C-terminal extension used to bind the portal clip [34]. Overall, the HT-adaptor is conserved across different phages with low sequence identity [17].

**Figure 5.**
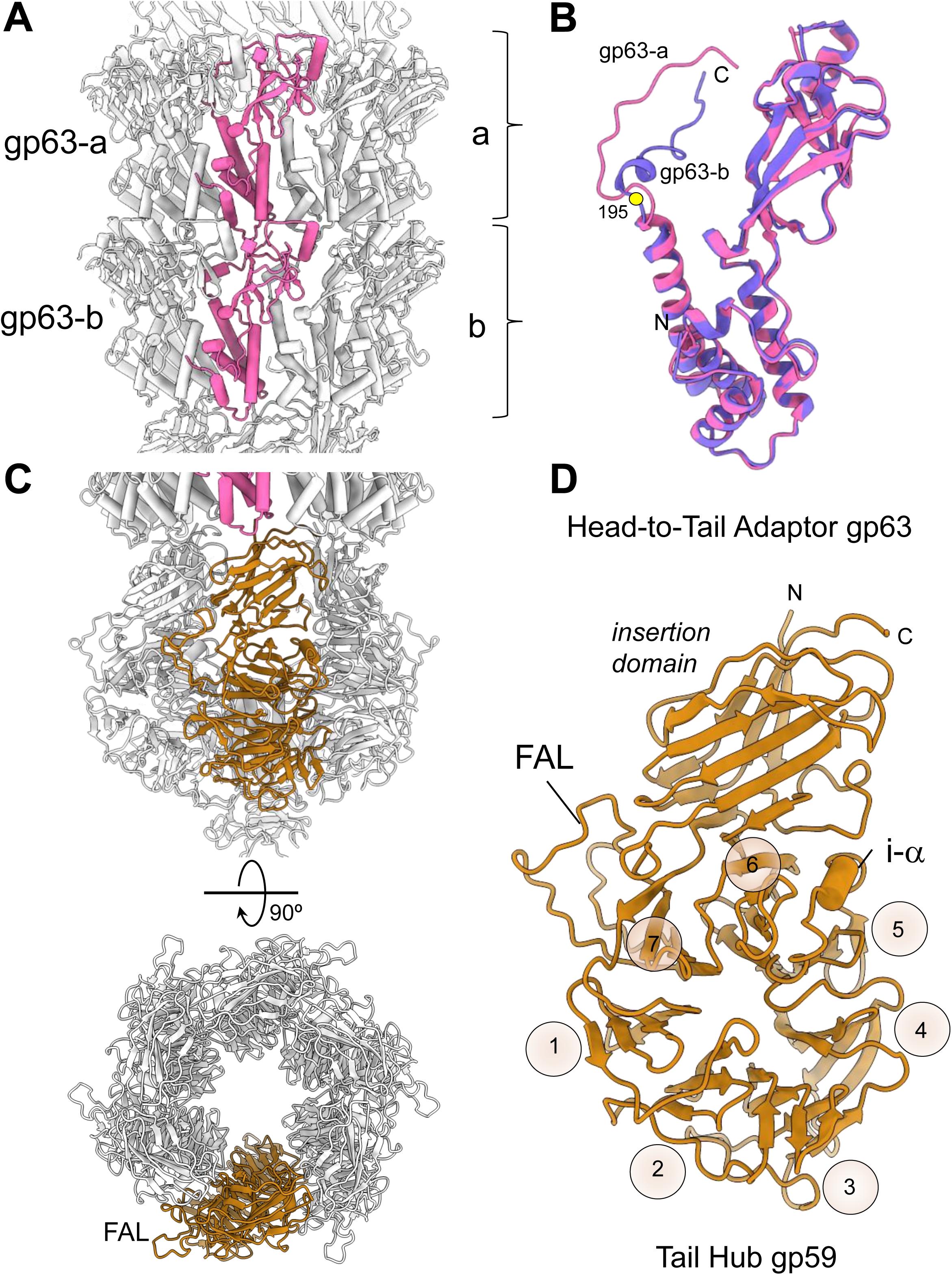
Structure of the tail proteins of *Pseudomonas* phage Pa223. (A) A side view of the Pa223 HT-adaptor shows two coaxial rings. In the diagram, only one protomer per ring is colored magenta, while the remaining eleven subunits are light gray. α-helices are shown as cylinders. The two dodecameric assemblies consist of the same subunit, gp63, which exists in two conformations: gp63-a and gp63-b. (B) Superimposition of gp63-a (magenta) and gp63-b (purple). A gray circle indicates residue 195, where the two conformers begin to deviate in structure. (C) Side view (top panel) and top view (bottom panel) of the Pa223 hexameric tail hub; only one protomer is colored orange, while the other five subunits are light gray. α-helices are represented as cylinders. FAL stands for Fiber Attachment Loop. (D) The ribbon diagram of the tail hub gp59 protomer reveals a six-bladed β-propeller. i-α refers to the insertion α-helix between blades 4 and 5.

The C6 symmetry tail map was used to build the Pa223 tail hub (gp59) and the tail fiber (gp60). The tail hub assembles as a 120 Å-wide hexamer, extending the channel formed by the portal:HT-adaptor complex by 85 Å and generating an internal 35 to 40 Å-wide lumen (Fig. 5C). We built a complete atomic model of the tail hub residues 2-513 that was real space refined to a CC = 0.89 at 2.8 Å resolution (Fig. 5D, Table 1) [41]. Pa223 tail hub protomer folds into an interrupted β-propeller, built by six four-stranded blades and an apical insertion domain. The insertion domain binds the HT-adaptor rings, whereas the β-propeller forms the tail nozzle. The insertion domain is formed by N-terminal residues 2-46 and C-terminal residues 403-512, while an extension of blade #7 (res. 354-387) forms the Fiber Attachment Loop (FAL) (Fig. 5D) used to bind gp60. FALs project outward in the hexamer (Fig. 5C) to recruit six trimeric fibers. Foldseek [33] identified the tail hub from the coliphage SU10 as the structure in the database with the highest similarity to the Pa223 tail hub (sequence identity 21.5% and RMSD = 12.7 Å), significantly higher than the T7 tail hub (sequence identity 10% and RMSD = 15.0 Å). The C6 map shows six tail hub protomers assembled into a hexameric quaternary structure with the individual β-propellors lying vertically, facing each other (Fig. 5C). Each tail hub:tail hub dimeric interface is stabilized by 28 hydrogen bonds and six salt bridges. PISA [38] estimates a ΔiG P-value = 0.882, indicative of minimal hydrophobicity and diagnostic of a weak oligomeric interface. Accordingly, phage P22 tail hub ortholog, gp10, ∼11.6% identical to Pa223 gp59, is monomeric in solution but oligomerizes only during assembly [42, 43].

### Pa223 Tail Fibers

Two types of fibers are visible in the asymmetric reconstruction of the Pa223 tail (Fig. 1A, B). Six tail fibers project upwards from the tail hub, as seen in T7-like phages [9, 44]; a single fiber, which has a weaker density, localizes at the tail tip distal from the portal protein, similar to the tail needle found in P22-phages [45]. To identify the gene encoding the major tail fibers, we scrutinized the C6 map. We observed excellent density for residues 2-123, which led us to identify this protein as gene product 60 (gp60) (Fig. 6A and 3B). We built these residues *de novo*, while residues 124-215, projecting upwards, could be modeled using AlfaFold2 and closely resemble the C-terminal tip of the phage T4 tail fiber gp37 (Fig. 6A) [46]. The entire tail fiber is relatively short, just 215 residues, a trimer of three identical subunits, was real-space refined against the 3.1 Å resolution, with excellent agreement between the model and experimental density (CC = 0.89) (Table 1). Pa223 fibers form a 45° angle with the 6-fold rotational axis that runs along the tail (Fig. 6A), but they also bend approximately 30° clockwise (Fig. 6B). Each trimeric tail fiber attaches to the tail hub through a single anchoring point, the FAL, that spans gp59 residues 354-387 (Fig. 6C). The 3:1 mismatched binding interface between tail fiber:FAL is dominated by hydrophobic contacts. Three tryptophan residues in FAL, W377, W370, and W363 generate a *pseudo*-three-fold symmetric surface that inserts perfectly at the base of the tail fiber (Fig. 6C). The tail fiber:FAL interface is likely flexible, and its flapping motion is the reason gp60 C-terminal tip (Fig. 6A) has smeared density (Fig. 1A) in the asymmetric reconstruction.

**Figure 6.**
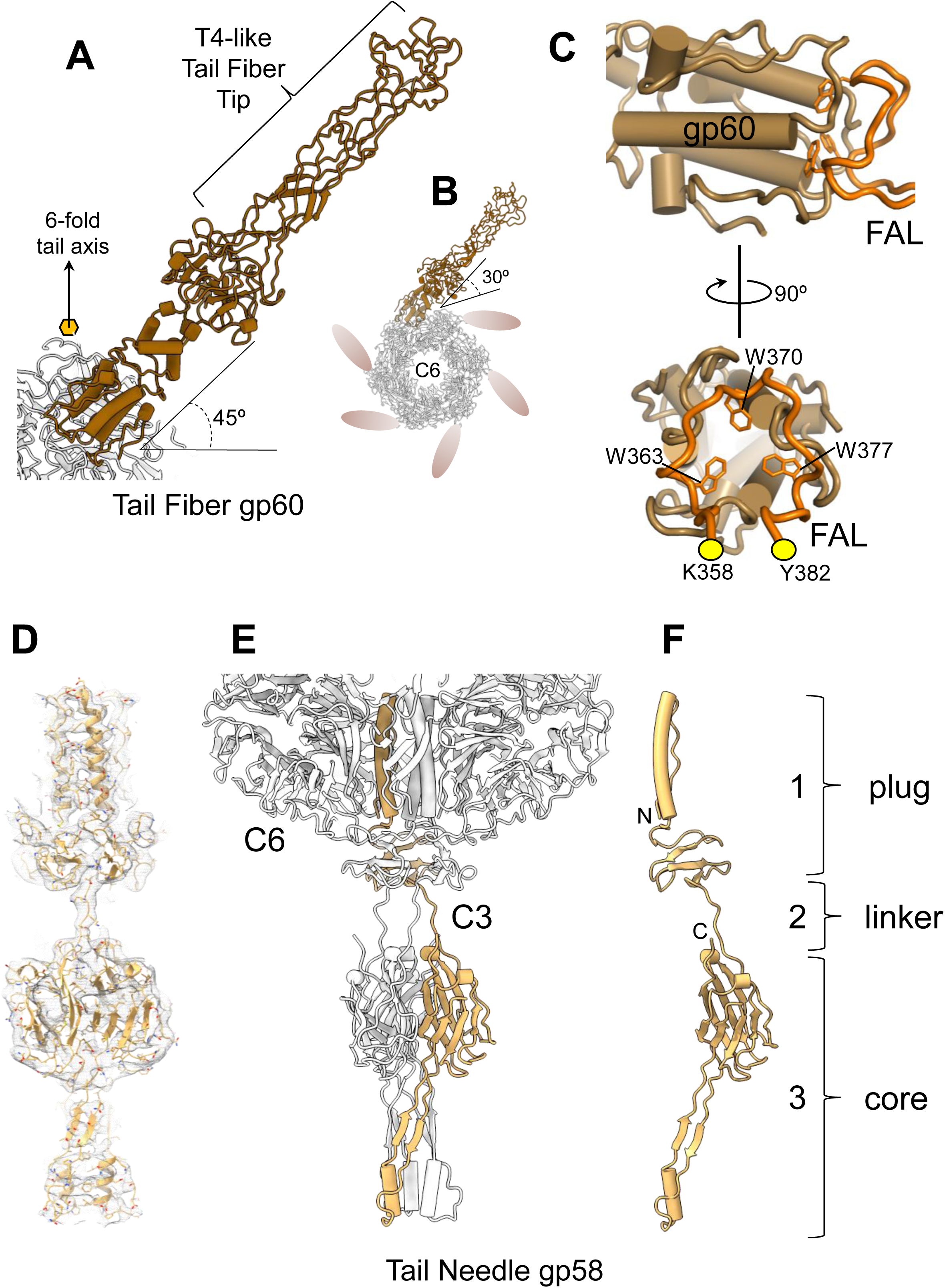
Pa223 tail fibers. (A) Ribbon diagram of Pa223 tail fiber (residues 1-340) colored maroon. The fiber forms a 45° angle with the six-fold rotational axis running along the tail. (B) Cartoon schematic of Pa223 tail hub (gray) visualized from above. Only one tail fiber is represented as a ribbon diagram, while the other five are depicted as maroon ovals. Each tail fiber extends outward in a clockwise direction and is tilted at a 30° angle relative to the tail. (C) A zoomed-in view of the interface between the N-terminal tip of the tail fiber gp60 (maroon) and the tail hub FAL (orange). Only FAL residues 358-382 are displayed, highlighting the hydrophobic core generated by W363, W370, and W377. (D) C3 averaged localized reconstruction of the tail needle gp58 calculated at 4.1 Å resolution and displayed at 3.0 σ. A ribbon model of the trimeric protein is overlaid with the cryo-EM density. (E) Ribbon diagram of the trimeric tail needle inserted into the tail hub lumen. Only one tail needle protomer is shown in light yellow, while the other subunits and the tail hub are depicted in light gray. (F) Ribbon diagram of the tail needle protomer. The brackets highlight three regions of the protein.

The density of the plug that closes the tail hub was weak in the asymmetric reconstruction of the entire virion, possibly underscoring the tail needle plasticity previously observed in similar protein plugs [47, 48]. We then computed a focused reconstruction of the tail tip that we improved by applying C3 rotational symmetry. The C3 localized reconstruction had complete density for the tail needle gp58 (Fig. 6D-F), which we built using a combination of *de novo* and Alpha fold-assisted prediction. The tail needle spans 150 Å, and the first 50 Å inserts inside the tail hub lumen, making several polar and electrostatic contacts through residues K4, E14, E16, E23, and R27. A flexible linker connects this plug to a trimeric core of mixed α/β fold, which likely moves relative to the tail, as suggested for the tail needle of phage P22 [48].

### Bioinformatic identification and structural modelling of Pa223 ejection proteins

The C12 reconstruction of the Pa223 portal revealed twelve copies of a helical protein bound around the portal crown (Fig. 7A) that we identified as gp54. This gene product is not a structural component of the capsid, but was also confirmed by MS analysis (Table 2). It lacks apparent homology to any protein of known function. Interestingly, a recent structure of the *Pseudomonas* phage DEV [45], a member of the *Schitoviridae* family, revealed a similar helical density surrounding the portal protein (Fig. 7B). Further studies identified the density around the DEV portal as the ejection protein gp72 [49], which is the ortholog of phage T7 periplasmic tunnel (PT) gp15 [50, 51]. Before genome ejection, gp72 forms a dodecameric assembly around the portal (Fig. 7B), which then folds into a nonameric tunnel into the bacterium’s periplasm after ejection. There is no evident structural similarity between Pa223 gp54 and DEV gp72 (Fig. 7C), which share approximately 6% sequence identity, undetectable through bioinformatics. However, both proteins are primarily α-helical and have comparable sizes (505 versus 521 amino acids). This led us to hypothesize that Pa223 contains ejection proteins within the capsid, similar to the larger *Litunavirus* DEV.

**Figure 7.**
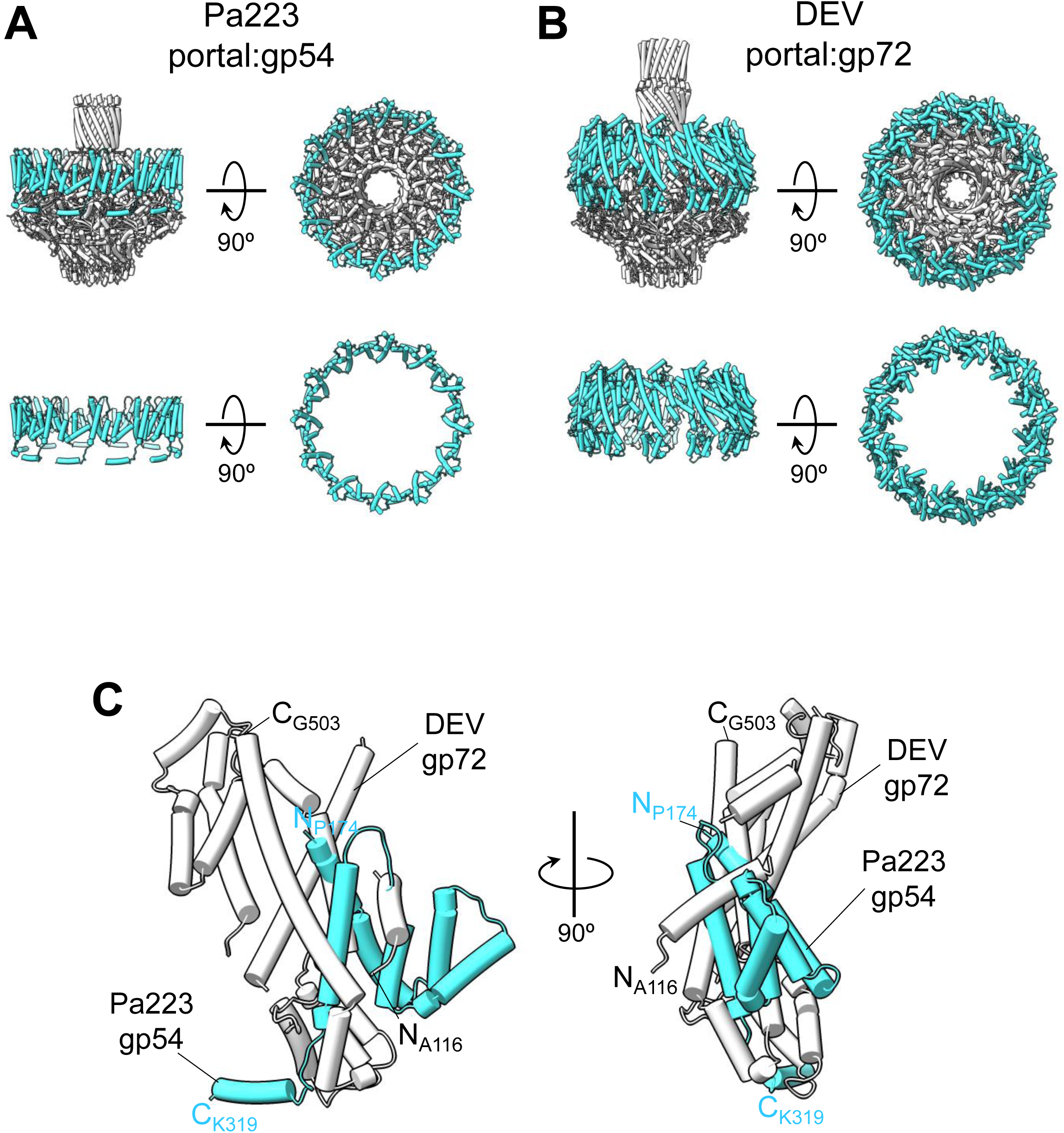
Similarity between Pa223 gp54 and phage DEV gp72. (A) Ribbon diagram of phage Pa223 portal (gray) bound to 12 gp54 (cyan) subunits. The bottom panel shows only the gp54 assembly. (B) Ribbon diagram of phage DEV portal protein (gray) bound to 12 gp72 (cyan) subunits. The bottom panel shows only the gp54 assembly. (C) Structural overlay of DEV gp72 (gray) and Pa223 gp54 obtained by superimposing the two portal proteins on which these factors are assembled.

If gp54 is Pa223 PT, we anticipate it to be flanked by two ORFs (Fig. 8A): one larger, about 1,000 amino acids, encoding the T7-gp16-like inner membrane complex (IMC), and the other significantly smaller, encoding a membrane protein that constitutes the outer membrane channel (OMC) of the DNA-ejectosome. A combination of bioinformatic and proteomic analyses confirmed this hypothesis. Firstly, MS analysis identified all three putative ejection proteins, gp53, gp54, and gp56, in the virion sample used for cryo-EM (Table 2). These gene products are similarly enriched and significantly above statistical significance, demonstrating that the three putative ejection proteins are virion components. However, only gp54 is visible in the cryo-EM reconstruction, likely due to its ordered conformation assembled around the portal perimeter. Secondly, the larger gene product 53 represents the putative gp16-like factor identified in T7-like bacteriophages, also called IMC [52]. This prediction was supported by a sequence alignment between Pa223 gp53 and T7 gp16 that revealed weak identity/similarity between these two proteins, possibly underscoring a common evolutionary origin (Fig. S3). Finally, the smaller putative ejection protein gp56 is anticipated to encode a membrane protein that perforates the outer membrane, similar to T7 gp14 [9, 50, 51] or DEV gp73 [45]. In support of this prediction, MemBrain [53] identified five transmembrane helices (TMH) in gp56, with three spanning the entire membrane bilayer (e.g., residues 233-243; 278-291 and 296-311) and two spanning only one leaflet (e.g., residues 80-89; 170-181) (Fig. S4). Interestingly, AlphaFold3 [54] predicts that gp56 folds into a hollow channel (Fig. 8B) only when the number of subunit copies is set to nine, as experimentally observed for DEV gp73 [45], which is a nonamer. Fewer copies result in a stack of gp56 subunits, instead of a hollow channel-like protein. Similarly, AlphaFold3 predicts that the putative PT gp56 assembles into a hollow channel (Fig. 8C) only when the search is conducted with nine copies. The shape of the predicted gp56 is highly suggestive of a periplasmic tunnel, with an overall length of ∼330 Å, an internal channel os ∼28 Å, enriched in basic residues, similar to previously visualized PTs, such as T7 gp15 [51, 55] and DEV gp72 [49]. For both gp56 and gp54 models, the confidence in the AlphaFold3 prediction is significantly stronger in the central region of the channel (e.g., Predicted Local Distance Difference Test, or pLDDT > 70) [54], which is involved in oligomerization, but decreases rapidly at the N- and C-termini, likely implicated in gp54:gp56 assembly and membrane insertion. For instance, phage DEV PT gp72 assembles onto the OMC gp73 by inserting the nonameric N-terminal end into the N-termini of gp73 [49]. Both N-terminal regions of the Pa223 AlphaFold3 models exhibit low confidence (Fig. 8B, C), suggesting that alternative foldings are possible. Finally, the function of the small gene product gp55 (Fig. 8A), consisting of only 55 amino acids and encoded between gp54 and gp56, remains unknown. AlphaFold [56] indicates that this protein folds into a continuous α-helix.

**Figure 8.**
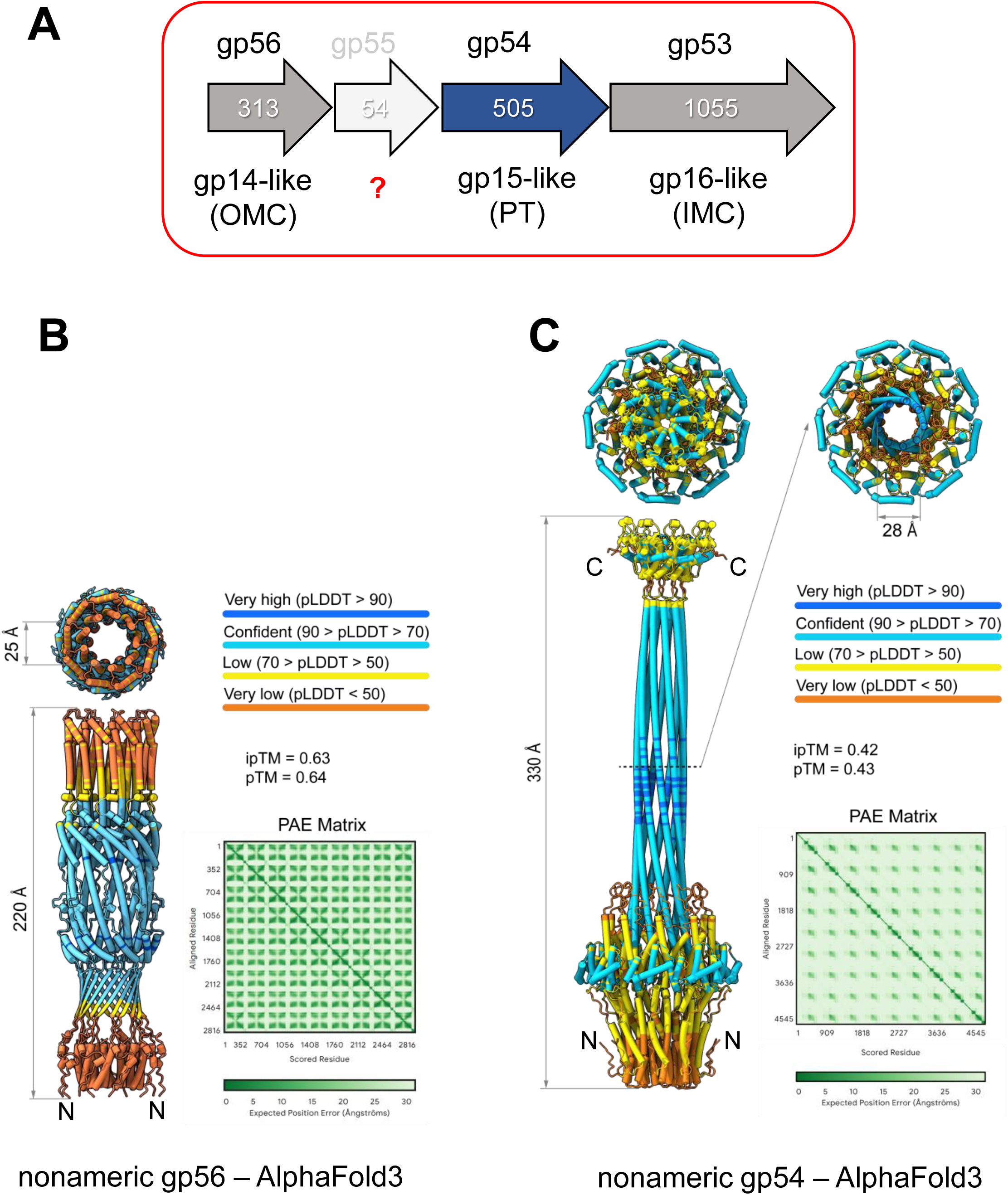
Identification of Pa223 ejection proteins. (A) Diagram of Pa223 putative ejection protein genes. OMC = outer membrane channel; PT = periplasmic tunnel; IMC = inner membrane complex. (B) AlphaFold3 model of nonameric gp56 (putative OMC). (C) AlphaFold3 model of gp54 assembled into a nonameric complex, possibly representing the post-ejection conformation of PT. Nonameric gp54 and gp56 are colored by AlphaFold’s Predicted Local Distance Difference Test (pLDDT), which estimates the local confidence in a predicted protein structure. AlphaFold3 server outputted statistics, including ipTM (Interface Predicted Template Modeling) and pTM (Predicted Template Modeling) scores, are included along with PAE (Predicted Aligned Error) Matrix.

## DISCUSSION

There is growing interest in *Pseudomonas* phages, which have potential clinical applications. Among these, phages of the *Bruynoghevirus* genus are attractive targets [8, 11, 12] due to their simple organization, broad host range, and potent lytic activity. In this study, we utilized an integrative structural approach to characterize the molecular composition of phage Pa223, one of the therapeutic viruses from Armata Pharmaceuticals’ collection. We obtained high-resolution density maps, sub-3 Å for most capsid and tail components, which allowed us to generate a comprehensive structural atlas of all Pa223’s structural proteins, and to identify and annotate the putative ejection proteins gp53, gp54, and gp56. This study highlights several aspects of Pa223’s biology that may have potential applications for other podophages used in clinical cocktails.

<u>First</u>, Pa223 and, by extension, members of the genus *Bruynoghevirus* share the same fundamental properties as those found in short-tailed phages that infect Gram-negative bacteria. These properties include the ability to recognize a bacterium through interactions with one or more receptors, bind to the bacterial cell surface, and assemble a channel through the host cell envelope to inject phage DNA, initiating a replicative lytic cycle. While these basic functions are conserved, the mechanisms and gene products that enable them vary greatly from genus to genus. Pa223 exhibits a uniquely mosaic composition characterized by structural elements previously observed in smaller T7-like and P22-like podophages, as well as in much larger *Litunavirus*. For instance, the tail of Pa223 has tail fibers that point upward, similar to those of T7, but they are considerably shorter and do not make contact with the capsid. At the functional level, the tail fibers of T7-like phages (*Teseptimavirus*) serve as primary receptor recognition devices to explore the bacterial surface, leading to a highly variable C-terminal domain that has been proven critical for host recognition [57–59]. In contrast, the tail fibers of *Bruynogheviruses* are highly conserved (90-100% protein identity), with no apparent hotspots of variability, typically associated with the need for creating receptor binding diversity. Therefore, it is questionable whether the tail fibers of *Bruynogheviruses* serve a similar function to those of *Teseptimaviruses*. Mutations in the hub or nozzle of other podoviruses have been shown to affect host recognition [60, 61], suggesting that the hub protein gp59 of Pa223 may serve as the primary host recognition protein. Alternatively, or in addition to the hub, Pa223 might use its tail needle as a receptor-binding moiety, similar to what is seen in P22 [62]. Consistent with this idea, the Pa223 tail hub does not seal like a nozzle, as seen in T7-like phages, but is instead closed off by a protein plug similar to P22 gp26 [63]. A conformational change caused by the interaction between the phage tail and the host lipopolysaccharide is likely to result in the ejection of the tail needle [64].

<u>Second</u>, a distinctive feature of Pa223 identified in this study, and likely conserved throughout the *Bruynoghevirus* genus, is the presence of two *quasi*-equivalent conformations of the HT-adaptor gp63, which forms two dodecameric rings that are coaxially assembled between the portal and tail hub. Notably, the *Gervaisevirus* phage GP4 [21] also contains two structurally homologous HT-adaptors encoded by two distinct gene products, gp81 and gp82. Multi-HT-adaptors are common in crassviruses [65, 66], where they are thought to lengthen the tail, providing more interaction sites for tail appendages.

<u>Third</u>, we identified a capsid spike, which decorates the 3-fold icosahedral axes and displays a triple β-helix that lacks obvious similarity to known structures according to Foldseek [33]. The striking feature of the Pa223 capsid spike is how the protein utilizes an asymmetric attachment domain to associate with the capsid, leaning 60° forward relative to the capsid floor. Pa223 is unlikely to function solely as a classical decoration protein [32], which serves as structural reinforcement for the capsid icosahedron. The superficial resemblance of the Pa223 capsid spike to a receptor-binding protein suggests that this protein may be involved in host recognition, potentially serving as an absorption moiety that prolongs the phage’s residence time near a bacterium, enabling the tail fibers, hub, or tail needle to identify their intended target. This idea is indirectly supported by the fact that Pa223 is one of the phages that are relatively difficult to separate from LPS during purification (*data not shown*). Every available genome of phages in the *Brunoghevirus* genus features a homolog to gp52, which strikingly displays slightly more variability than the tail fiber proteins (87-99% identical), suggesting higher selective pressures for their diversification. Admittedly, little is known about the biology of *Bruynogheviruses* to speculate about the exact function of the capsid spike, which future structure-function studies will need to investigate.

Lastly, our study led us to identify Pa223 ejection proteins, which seem somewhat different from those previously characterized in other phages [52]. To date, three distinct arrangements of ejection proteins have been described before genome ejection in short-tailed phages. In phage T7 [9, 10, 67], and possibly in all T7-like phages, the ejection proteins form coaxially stacked rings on the portal protein, with stoichiometries consistent with four copies of the largest subunit gp16, eight copies of gp15, and eight copies of the smallest subunit gp14. Importantly, portal proteins from T7-like phages lack a barrel domain [68], which allows ejection protein oligomers to stack upon each other and the portal. After infection, the ejection proteins reorganize into a trans-envelope channel or DNA-ejectosome [69], where gp14 and gp16 span the outer and inner membranes, respectively, while gp15 forms a PT connecting the two membranes [51, 55]. In contrast, phages with a portal protein whose C-terminal domain folds into a barrel, such as P22 [19] and Sf6 [34], cannot accommodate stacks of ejection proteins arranged coaxially onto the portal, unlike T7. As a result, they have ejection proteins that are probably loosely associated with the portal barrel [70]. Cryo-EM studies of P22 mutants lacking individual ejection proteins [19] revealed that the barrel unfolds without these proteins, supporting the idea that the ejection proteins interact with and stabilize the barrel. Furthermore, the genetic deletion of the ejection proteins from the virion causes P22 to package a DNA molecule longer than average [71], indicating that the ejection proteins occupy space within the virion that can be filled by packaged DNA when they are absent. However, the ejection proteins are not visible in P22-like phages through cryo-EM SPA because of the technical limitations in aligning these proteins within the DNA-filled capsid. Despite differences in location and assembly within the capsid before ejection, after ejection, the P22 ejection proteins form a transenvelope channel similar to phage T7 [72], suggesting a comparable function. Finally, recent work on the *Litunavirus* DEV [49] revealed an ejection protein surrounding the portal perimeter. This feature was also observed in the phage ΦcrAss001, where ejection proteins are described as cargo proteins ready for ejection [66]. This discovery led to the identification in phage DEV of an operon encoding three ejection proteins, which include a short gp14-like membrane protein, a gp15-like PT nonameric after rejection, and a giant gp16-like protein, approximately 3,500 residues long, which functions as virion-associated RNA polymerase (vRNAP) [73]. The ejection machinery of Pa223 appears to be conserved in its genetic synteny across the *Bruynoghevirus* genus, although the genes themselves can diverge notably in sequence, dropping to as low as 83.8% protein identity for gp54. In this study, we describe the *Bruynoghevirus* ejection proteins, which exhibit hybrid features that are distinct from the three groups mentioned above. Similar to P22-like phages, the Pa223 ejection proteins cannot stack onto the portal protein due to a prominent barrel domain that extends 150 Å into the capsid. Unlike P22, the gp15-like subunit gp54 assembles concentrically around the portal perimeter, like the much larger *Litunavirus* DEV. Upstream of gp54, Pa223 features a gp14-like membrane protein, gp56, which is roughly twice the size of its counterparts in P22, T7, and *Litunavirus* phages (313 vs. ∼150 residues). Downstream of gp54 is the larger gp16-like subunit gp53, comprising 1,055 residues, which shares limited similarity with the N-terminus of N4 vRNAP, a protein also known to be ejected into the host [73]. Finally, Pa223 has a short ORF between gp54 and gp56 that encodes a predicted 55-amino acid protein, the function of which remains unknown.

In summary, this study expands the repertoire of *Pseudomonas* phages resolved at near-atomic resolution, providing valuable insights to understand variations in phage specificity, stability, and resistance mechanisms. The 3D atlas of Pa223 structural proteins presented in this paper will help map mutations that alter phage functionality and facilitate the targeted optimization of phages for therapeutic applications.

## MATERIALS AND METHODS

### Origin and characteristics of Pa223

Pa223 was isolated from sewer samples of the greater Sydney metropolitan area, Australia. Pa223 was part of a phage cocktail candidate developed by Armata Pharmaceuticals. Fermentation and purification were performed using proprietary methods in order to achieve clinical levels of purity and a titer of 1 x 10^13^ PFU/ml. The current annotations for Pa223 were generated from Pa223’s published sequence (accession MK837012.1). Glimmer was used to identify additional putative protein-coding sequences and alternative start sites [74]. ARAGORN was used to identify tRNA and tmRNA genes [75]. Sequence alignments of *Bruynogheviruses* from public databases and Armata’s collection were conducted using MAFFT [76] and manually curated to resolve ambiguities with the assumption that ORFs or start codons present in the majority of the phages are more likely to be real than those present in fewer members of the genus. Finally, CDSs were renamed to ensure consistency in their names across members of the phage genus, making future comparisons easier. A list of putative coding sequences is provided in **Table S1**.

### Vitrification and data collection

2.0 µL of virions, concentrated at low-speed approximately 10-fold starting from a PFU of 1 x 10^13^ phages/mL, was applied to a 200-mesh copper Quantifoil R 2/1 holey carbon grid (EMS) previously glow-discharged for 60 sec at 15 mA using an easiGlow (PELCO). The grid was blotted for 7.5 sec at blot force 2 and vitrified immediately in liquid ethane using a Vitrobot Mark IV (Thermo Scientific). Cryo-grids were screened on a 200 kV Glacios (Thermo Scientific) equipped with a Falcon 4 detector (Thermo Scientific) at Thomas Jefferson University. EPU software (Thermo Scientific) was used for data collection using accurate positioning mode. For high-resolution data collection of the Pa223, micrographs were collected on a Titan Krios (Thermo Scientific) microscope operated at 300 kV equipped with Falcon 4 direct detector available at Stanford-SLAC CryoEM Center (S2C2).

### Liquid chromatography/mass spectrometry (LC-MS/MS) analysis

Phage samples were treated with 12 mM sodium lauryl sarcosine, 0.5% sodium deoxycholate, and 50 mM triethyl ammonium bicarbonate (TEAB), heated to 95 °C for 10 min and then sonicated for 10 min, followed by addition 5 mM tris(2-carboxyethyl) phosphine and 10 mM chloroacetamide to reduce, and alkylate the proteins in sample. The sample was then subjected to trypsin digestion overnight (1:100 w/w trypsin added two times). Following digestion, the sample was acidified, lyophilized, and then desalted before injection onto a laser-pulled nanobore C18 column with 1.8 μm beads. This was followed by ionization through a hybrid quadrupole-Orbitrap mass spectrometer. The most abundant proteins were identified by searching the experimental data against a phage protein database, the *Pseudomonas* host protein database, and a common contaminant database using the MASCOT algorithm [77].

### Cryo-EM SPA

*Pseudomonas* phage Pa223 micrographs were motion-corrected with MotionCorr2 [78]. RELION’s implementation of motion correction was applied to the micrographs with options of dose-weighted averaged micrographs and the sum of non-dose-weighted power spectra every 4 e^-^/Å^2^. CTF (Contrast Transfer Function) was estimated using CTFFIND4 [79]. After initial reference picking and 2D classification, the bin to 4 particles were subjected to a reference-free low-resolution reconstruction without imposing symmetry. The particles were then 3D classified into four classes, with I4 symmetry imposed. Of the four classes, the best class was chosen and was subjected to 3D auto-refinement to align the particles finely. The particles were then expanded according to I4 symmetry using RELION’s *relion_particle_symmetry_expand* function to obtain 60 times the initial particles. A cylindrical mask (r = 200 Å) was generated using SCIPION 3.0 [80] and then resampled onto a reference map covering the five-fold vertex in UCSF Chimera [81]. The cylindrical mask was then used for non-sampling 3D classification (as implemented in RELION 3.1.2 [82, 83]) without imposing symmetry to search for the tail.

Locally aligned particles were then combined, and duplicate particles were removed. The particles of correct tail orientation were then re-extracted by shifting 300 Å along the z-axis with an unbinned 512 pixel box. The initial localized reference map was reconstructed directly from one of the classes using RELION’s *ab initio* 3D Initial Model. Selected 3D classes were auto-refined using C5 symmetry, followed by five-fold particle expansion. The expanded particles were subjected to a third 3D classification without sampling, and the map was symmetrized by imposing C12 and C6 symmetries, which gave the best density for the ejection protein, portal, head-to-tail: tail hub, tail fiber, tail needle complexes, respectively. Furthermore, a focused, tight mask was used to reconstruct the tail fiber without applying symmetry. Reconstruction of the capsid spike was done by realigning the capsid particle from I4 to I1, locally re-extracting the 2-fold vertex by moving the box along the z-axis to the center of the capsid shell, where it covers two capsid spikes. Then, a focused mask was applied to one of the capsid spikes, followed by non-sampling 3D classification to identify an asymmetric unit of the trimeric capsid spike. All steps of SPA, including 2D Classification, 3D classification, 3D refinement, CTF refinement, particle polishing, post-processing, and local resolution calculation, were carried out using RELION 3.1.2 [82, 83]. The final densities were sharpened using *phenix.autosharpen* [84]. RELION_*postprocess* [82, 83] was used for local resolution estimation. All cryo-EM data collection statistics are in **Table 1**.

### *De novo* model building, oligomer generation, and refinement

All *de novo* atomic models presented in this paper were built using Coot [85] or UCSF ChimeraX [81]. We used the following cryo-EM maps for model building: (***i***) A 4.0 Å asymmetric reconstruction of the entire Pa223 virion. (***ii***) A 2.6 Å C5-averaged localized reconstruction of the capsid revealed the atomic structure of the Pa223 coat protein (res. 1-317). (***iii***) A 2.6 Å C12-averaged reconstruction of the unique vertex was used to build a model of dodecameric portal protein (res. 2-639) bound to two dodecameric HT-adaptor rings (res. 2-208). The portal protein has no visible density of residues 1-10 and 640-706 in the C12 averaged map. The ejection protein gp54 was built in a 3.0 Å focused reconstruction of the 2.6 Å C12 map obtained using a smaller mask. (***iv***) A 2.8 Å C6-averaged reconstruction of the tail was used to identify and model tail hub gp59 (res. 1-513). The tail fiber gp60 (res. 1-215) was modelled into a C1-focused reconstruction of the tail fiber at 3.1 Å resolution. (***v***) A 3.0 Å C1-focused reconstruction of the capsid spike was used to identify and model gp65 residues 2-317. (***vi***) A 4.1 Å C3-averaged local reconstruction was used to model the gp58 tail needle (res. 1-208). All atomic models were refined using several rounds of rigid-body, real-space, and B-factor refinement using *phenix.real_space_refinement* [86] and validated using MolProbity [87]. Refinement statistics are in **Table 1**. The tail fiber gp60 and tail needle gp58 prediction models were generated using AlphaFold [56] and AlphaFold2 [88]. Oligomeric gp54 and gp56 were predicted using AlphaFold3 [54]

### Structural analysis

All ribbon and surface representations were generated using ChimeraX [89]. Drawings of electron density maps and local resolution maps were generated using ChimeraX [89]. *De novo* density building was attempted using ModelAngelo [90]. Structural neighbors and flexible regions were identified using Foldseek [33]. Binding interfaces were analyzed with PISA [38] to determine bonding interactions, interatomic distances, and types of bonds. Membrane insertion was predicted using MemBrain [53]. All RMSDs in the Cα position between superimposed structures were calculated using Coot [85].

## Abbreviations used

cryo-EM: cryogenic electron microscopy
LC-MS/MS: liquid chromatography/mass spectrometry
HT-adaptor: head-to-tail adaptor
M.V.: mature virion
PT: periplasmic tunnel
OMC: outer membrane channel
IMC: inner membrane complex
SPA: single-particle analysis
CC: correlation coefficient
RMSD: root-mean-square deviation
dsDNA: double-stranded DNA
SSM: secondary structure superimposition

## ACKNOWLEDGMENTS

This work was supported by National Institutes of Health grants R01 GM100888 and R35 GM140733 to G.C. A portion of this work was carried out at Stanford-SLAC CryoEM Center (S2C2), which is supported by the NIH Common Fund Transformative High-Resolution Cryo-Electron Microscopy program (U24 GM129541). The mass spectrometry work was conducted under contract by the Pasarow Mass Spectrometry Laboratory at the Semel Institute for Neuroscience and Human Behavior at the University of California, Los Angeles.

## AUTHOR CONTRIBUTIONS STATEMENT

C-F.D.H. and G.C. performed all steps of the cryo-EM data collection and analysis, deposition of atomic coordinates, and maps. G.C., P.K., D.B., and S.L. supervised the entire project. C-F.D.H. and G.C. wrote the paper. N.B. and R.K.L. aided in data analysis, interpretation, and making figures. S.B., J.R., R.G., A.S., and L.S. amplified and purified Pa223 for cryo-EM analysis. A.S. analyzed the LC-MS/MS data. R.G. and S.L. sequenced and analyzed the genome of Pa223. All authors contributed to the writing and editing of the manuscript. The atomic models and three-dimensional reconstructions described in this paper are available in the Protein Data Bank (9NY2, 9NWI, 9NXK, 9NWM, 9NXO, 9NY6, 9NXP) and Electron Microscopy Data Bank (49916, 49926, 49882, 49902, 49887, 49909, 49925, 49910), respectively. All other data are available from the corresponding author upon reasonable request.

## COMPETING INTERESTS STATEMENT

J.R., R.G., S.B., A.S., L.S., P.K., D.B., and S.L. are employees of Armata Pharmaceuticals Inc., a company involved in the development of bacteriophage therapies. The other authors declare that the research was conducted in a way that is free of financial or commercial relationship that could be construed as conflict of interest.

## Notes

### Competing Interest Statement

The authors have declared no competing interest.

