## Supplementary material for "High-resolution cryo-EM analysis of the therapeutic *Pseudomonas* phage Pa223": Suppl Fig 1-4 and Table S1

***Corresponding Author:**

Gino Cingolani, Ph.D.

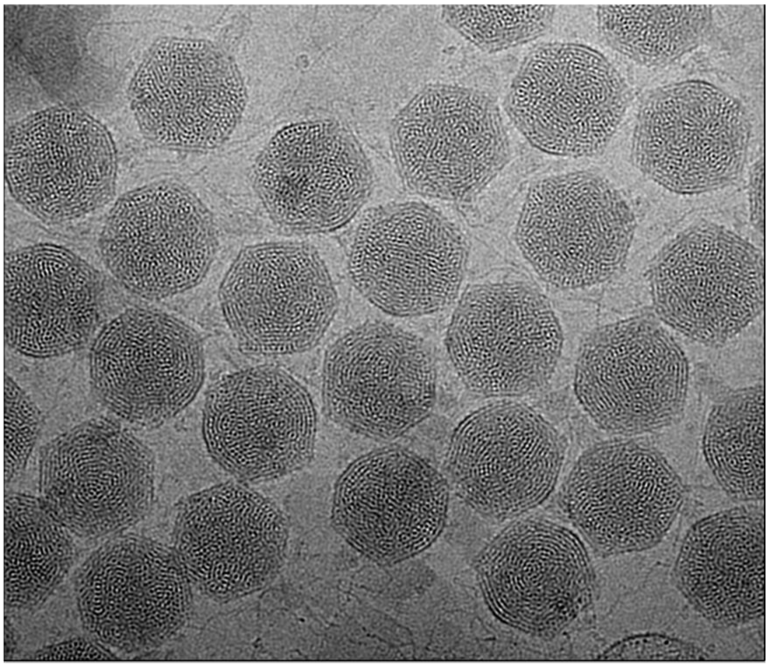

B

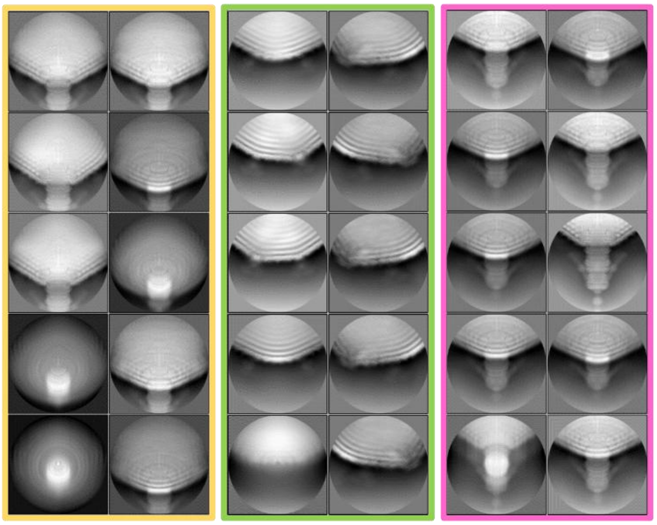


**Figure S1. Cryo-EM analysis of phage Pa223.** (**A**) Representative cryo-micrograph of purified Pa223 virions. (**B**) Representative localized reconstruction of 2D classes of the capsid protein (left), capsid spike (center), and tail apparatus (right).

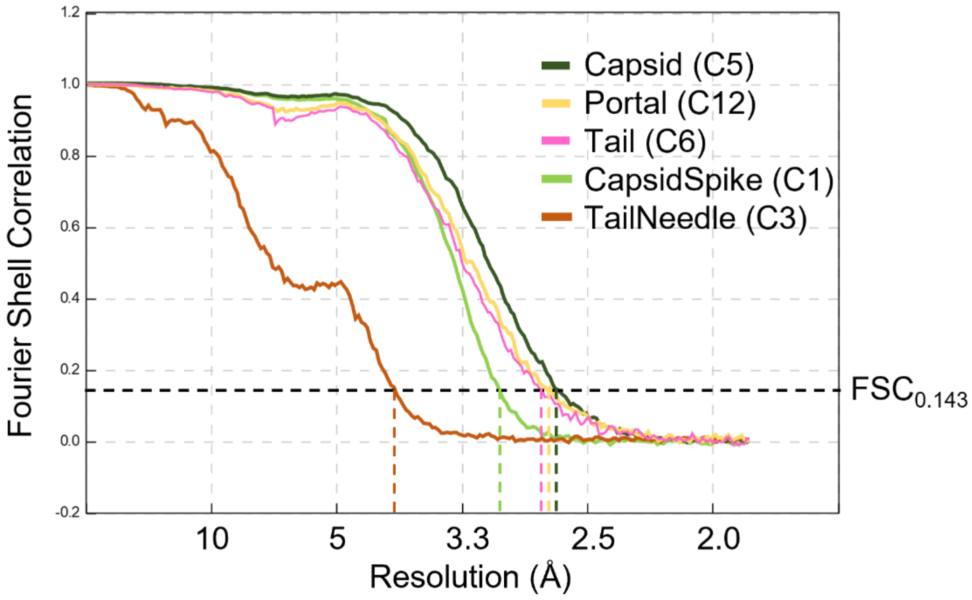


**Figure S2. Fourier Shell Correlation (FSC) resolution curves** for all reconstructions presented in this study. For all curves, the resolution is indicated at 0.143 cut-off.

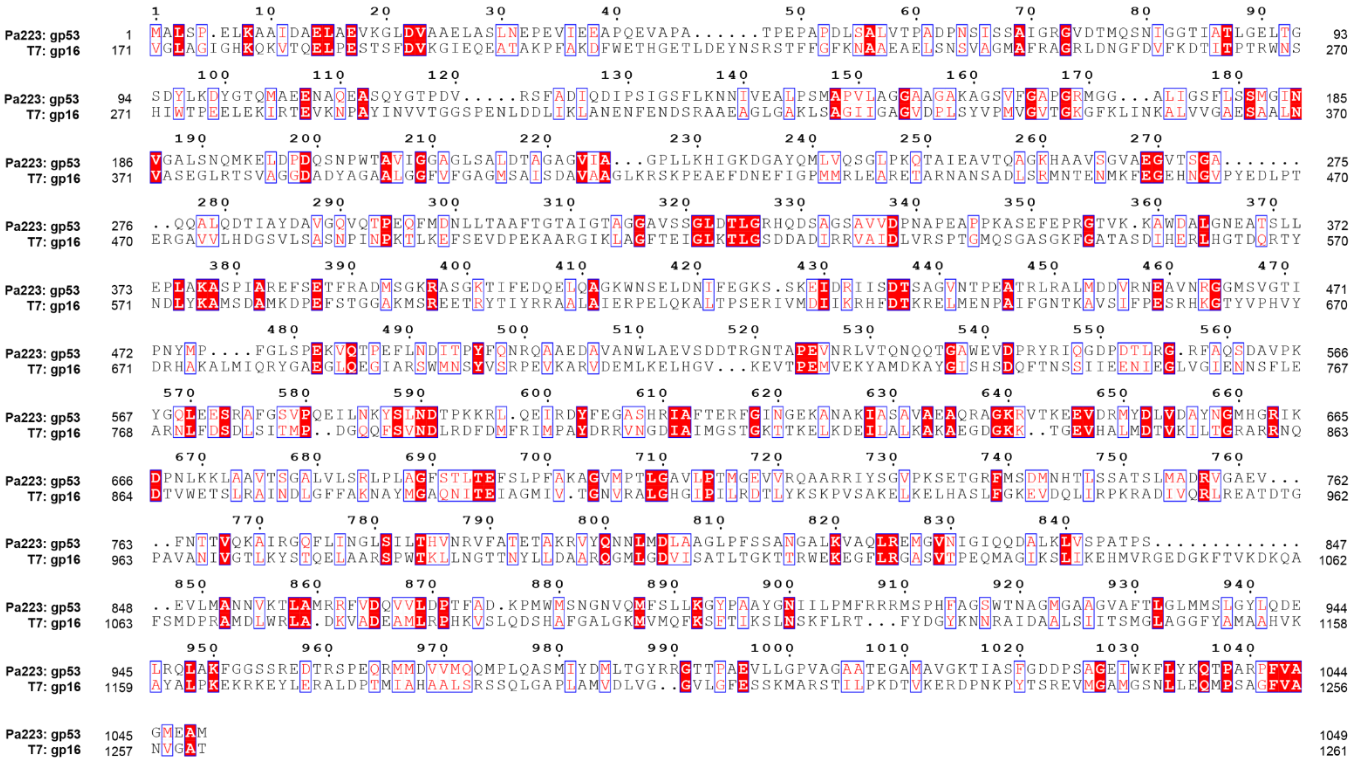


**Figure S3: Sequence Alignement T7-gp16 vs Pa223-gp53.** ClustalW alignment of T7 gp16 and Pa223 gp53 represented using Esprit. Identical residues are boxed in red with white text, and similar residues are boxed in white with red text.

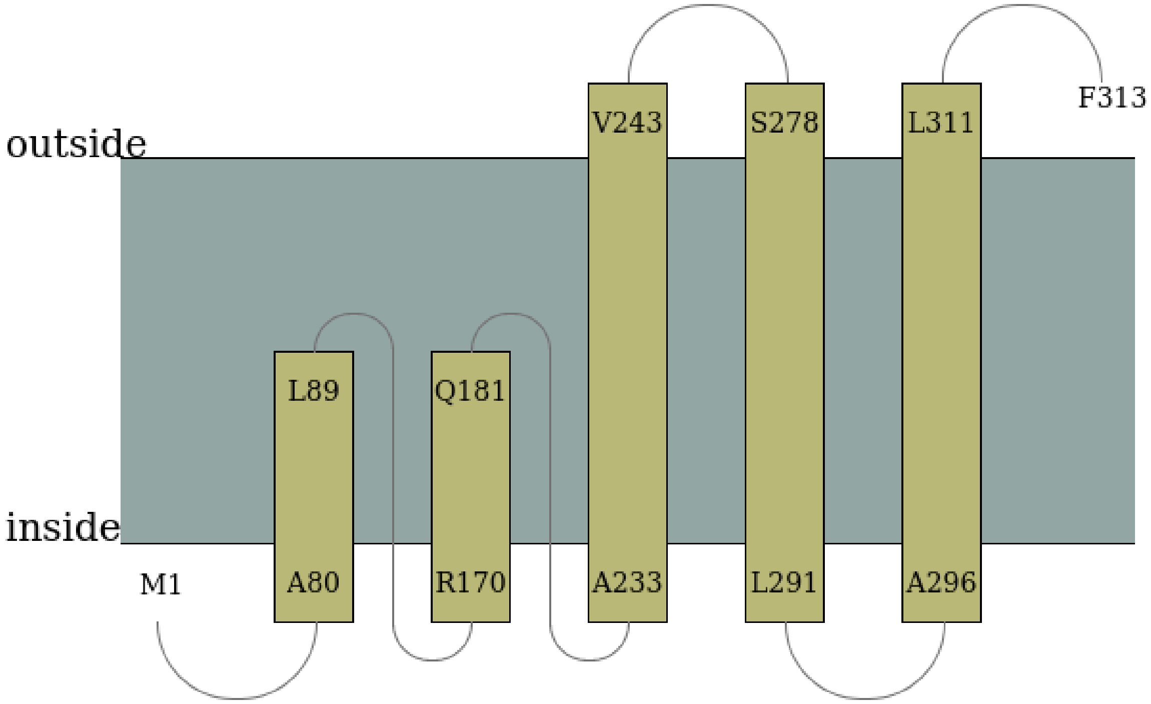


**Figure S4: Cartoon representation of gp56 topology predicted by MemBrain.**

**Table S1: List of predicted proteins encoded in the genome of Pa223.**

| Name | Length | Sequence | Function | Source |
|---|---|---|---|---|
| gp01 | 102 | MNAALDKKLRRKASRKARAQGLNIGHLGKAQQRAEQKFGIISNCNKILDSKFASLQEKAGARKRKALMNTDWRNREDTKLRNWYKPSKCKNSPVTHIKPEV | | |
| gp01-1 | 46 | MAKVGARDSKTPEDSQRTDHNEDTNAERGRLRQCQGLPTNPFIVS | | |
| gp01-2 | 61 | MGKCTFDGPWTEQGRNPKAGGITHTPEERRFWPCTIEGPTGCLLQRKQLAEFPVESWGTV | | |
| gp01-3 | 29 | MATQKKSLCRRTGRTNPRQRVDKPSPRL | | |
| gp01-4 | 31 | MHDSLGISVALTSKTDGHSEKEPMPENRTN | | |
| gp01-5 | 65 | MTNPAEPGLMATTFHLGHHPPSLRTLTGAANQRGMPKSPKYKTGRHRGLTRDRGSGKMAPKSTA | | |
| gp02 | 161 | MSDYKRINGIIKTIAQRGASLDKLIQTTGLDILKHIEEHGEVSLACKLFKAMPQGSRRNALAAWFIDHGKIEANIDKEKSKEFPFVIAKDKATRLERAAEKPWFKYKKERDVADEFSLDQAIAAFKAKIQRAIDKGQLQAADERIAVIQRLEVKDEAKAA | | |
| gp02.1 | 24 | MYYDAPLGSSLEYMVGPCIVWMN | | |
| gp03 | 73 | MDELTEDKTVWTPFGSFLVAVTNDYVQGRCNSRSYNSDGSEQVVYRWAFVDGKFACGVYVDGRPMGFCGWTK | | |
| gp04 | 47 | MKSTYEAAHERALMVNRLQKLTRMLRVHHDPKWKQEQQELIKRLRR | Homologous to Mip | Wagemans et al., 2015 |
| gp05 | 42 | METGTTGTYQEVKAMIISIVVSCVGIGYFFFRDWKEEMGIN | | |
| gp06 | 49 | MQAIEAIDRKIAKVTRELTNSSGNHLGKTLELNKLRAKRASLAKNRAR | | |
| gp06.1 | 42 | VMSWKYIACAAIRSRSNGLLLVAWVVLFVLSSTLDSIPFAS | | |
| gp07 | 78 | MKINVVEKPKKVPAHTLKATDVFRVENIHGMVLRDDHNHYLPNDGDVYFVCLEAGIICSIHSDCEVIPVDAQMTVRV | | |
| gp08 | 52 | MTPALYLLLWLESLVSIRRFQPNHKAYPGWYVTHIRVSIFGKRVGIVYEIH | | |
| gp08.1 | 49 | MKSTKPQVFWLVSKDGERHVKVINFGQFITETLYELGIPVDHIVHRVQ | Homologous to Igy | De Smet et al., 2021 |
| gp09 | 88 | MTNTSFTVAVAKPRFEFHISKMADGAYVHSGDSGSVDLVYLKKGELVTIIHKDGFNILKCKEIGTHPDLFPAKWLSVNACMWGPYEE | | |

181

| Name | Length | Sequence | Function | Source |
|---|---|---|---|---|
| gp10 | 89 | MIGQIMAQERRMKRRVEKRGFNMSLQEGTQEPRGELGFTLAAVGMESSRSAYLRHAREAMIQSGEPCPHC MPIFGPKEGRCCNCARDW | | |
| gp11 | 91 | MGRSKHLTFLPSRITVLDDDTVYVPWRYQLESLYNMASAGGATDPEAVKGEFLDWVRAEYGRYVSIELEGIPE DREDAFDEYVLKGSQQG | | |
| gp12 | 85 | MMTYLLIIPAILIYMAFSLLVAGIAGLAANCDEYGRMSQKDQDVSIILGILWPVSLPWMCFMCVIWNPLATTIR AAKRLIKGDY | | |
| gp13 | 142 | MRRIDFPSCCTAKIYIGMGPSGTADHRAGEASNGFSPRGFAKELIGVIRREYNEGHGTAVFSVNNEQVVADTIL RRMGSHYNPWASSYNHSTKVRVHVINVQAAAEILINHGVLIRRLGGLQDYPGTVEHSEYLDKLCKGL | | |
| gp14 | 80 | MVLKLYTRVMLLAIPAWISILWVVPQMLSPADTMVNAIGAALLVISILYSLWAVSHLYSYASKLSGKKEVHRGH GNEKQ | | |
| gp15 | 73 | MATVTRSSKVPTSPLYFERKCDSQIYLRSESRKDLTIPMVNICNGSVLYLHPRELPVYFVAIPEGETLKVKV | | |
| gp16 | 73 | MTEQQRDRLRKAIHMDLEGIGEDTPDLYDNIHKIEGLLGHPGKPPHDWGFYVLMFVMFISLCMVYITPLLYK | | |
| gp17 | 275 | MKKILGLLAIAAIAITAGCSKVPAGYTGVVVNMMGDSKGVDLKETPVGWKFLTPNEELFKFPTFNQNFDLAVV TAQDKDGLKLDFPIGVTLRAAPGSAPLLFQTYRKGMNEIVSVNVPQVVRNAVNNASSKKSAEAIYGPGKEAFV KEVEAAVREHFSTRGIIVESLYLNGMIGLPAQVVEAINAKIKATQTAMQRENELRQTQAEAAKAIAAAEGEKQ AAILVAQGKAESLRIQGEALRQNPGVVELNAIEKWDGKLPTYVTGGQAMPFINVR | | |
| gp18 | 43 | MGKRKVFYERICSNGELTLNGNLKATKVSAPTKGKQQRRGPF | | |
| gp19 | 321 | MARIKYAFGMKSKKGGVKALKVMTSASCFGAMEGVVTHGYKLDGWTFVCSRRSKKFIDALNKCTQGELKTIT IGGKAYKIPKVHFWSYESKAKTSPFVKYVSGSELISPTQHGKEGVCGIYFNPKEHTLDSWYPIMKFLFKLVSSGLD PQGREEKIHMELAEKYGFWKSYLAMSFHGLVANGYTGYPLTNYMFSNDWEDIKKGDIHIQSAEECNRFGRW VPNRNAPGGREWVRSKPYKSEYLGIKLNKVEVVTIYPQPTIFGKKTPYDIVDTELGGFLRLTPSDCEKHGVVML GTMGEIHWGRGSRDIPLALEGFIEKNF | | |
| gp20 | 319 | MGMYAAHNVYYDAEGAEVGHCCVKYVAEYTNCFGAFCDSTGPHPDIVWETGVGHLLVSTHENTAGKLVEFL NSDLVNRITDGGILSASQDWPTEWWTGSGRKQNLSNPDHFSPPGRRQAVYVRVDLKKNASAIICALRMGDR LWGIGDQMRRIPKEAQDKILGFNAELLTLAACAQVGSTAHCDSYPAMFPVTAGEYKEQCEANGDDVEDSYIS EIMDHLTGGLKDITYFDIKSIHGRTREEFKDKLKHHNDPLWLGYTKDDYLIECDGLDGVHDDRILSIMAPITIDV GDPGRDNSPTVPTGYYDLPEILEKLESMQ | | |

| Name | Length | Sequence | Function | Source |
|---|---|---|---|---|
| gp21 | 267 | MNNAIPLIGADPEVFVGYDRNPQSVIGFIGGTKEAPLAVAGGAVQEDNVLLEYNIDPASTKEDFVKRIISVRLLG<br>AQMLHPFGMNIIENLSSHLYDEELLRSFGPQAYVFGCEPDYNCWTRRQNVMPNDAPPTLRTAGGHVHIGFG<br>HIERVTKATTREVMQMCDYLLGLASVLLDGDTQRKKLYGKAGAMRYKPYGGEYRSLSNFWIFSVNLTEWVYE<br>MAVQAYTSKHLLEEYKSIVSGDEVQRIINENDGVAAVAALQALGVKYE | | |
| gp22 | 190 | MNDLNNRHRLAGDFNMYYSSTFAFFRVDGEPRVVYVDDTESIGDDRQFDGFRLLGNIFRPDGGQYYGGVVY<br>SEVEGVRPPSGYYDVFGGGERDTYVSFLVNNRTQRKGVDPRNVLINHGQQAVTGQMMIRIFLQAEEMISDP<br>AHRDFFIKDGVVNWKGVKVGQMVDNRLSVDEQFKNQEDLLCRLLARR | | |
| gp23 | 392 | MSVIGTQIGFRKNQIKTPEHHEELPAVASFGFEVELEGLDQWPEVDGWDIKNDGSLRNGMEYVFSGPASGE<br>QAINRVEAFANAMEQNPPAPTFRCSTHLHMDMRDVNWKVYERTVLAYMVFEDVFFDHCQPYRRDSNFCIP<br>FFSNDWLSQTFGRRILAPEGDREKVLGLTSWPKYSALNLQVTHNFGSIEFRGAHAMTTRSEMVGLMQRMM<br>CLKAFAIAHEETPLGEFVKVLSDVNLRDIFFLGVAPDYEMSPGGREMGIASASLAIATMGFVRSGVDPLEDEQN<br>RQRRLREQAREQQRALDRRLMAARSVLGRIREGAPERYNLAIIEGSNMRMDQVLTTVQSLRMIGHQVSVRD<br>LVEDNEILRDSFVLLLDNPDHMQRQAGFRLEENMY | | |
| gp24 | 511 | MCGLVGFCATTNASDNEIALLKSLLAVDIIRGAHATGLAKIDPVKNEVGIHKRAVDAYDFLADPETKEFLDKGR<br>ARIYMGHNRYATMGDKTDHANAHPFQVDHITMVHNGTVDSWGLHLLDGNDKYNVDSNMLCATIANHGA<br>KKTFEEKFSGAAAVIWWDSKERTLNFIRNDERPLFMAVTTTGTIVWASEPGMLKVFLERPNAKIRLRSPIAELK<br>AEVLVTIPFTEAGVRKGVEPQTTPVTFLDLPIPESERQAAAWWSRYVGVSDYEDYSRSQGSQAGTKGSKAGSS<br>YGTSGDAYARNTLRINNNLDAAGSTFKHRQLVTFDVVKVEAYANGSEYGTVTGIEREENLLIEAHGINVAKVH<br>GYTVLRGSISNAYFIGQDRDLKVTVEDLAVSCLDPKHRPTPGETTPVLRIGTISSETKSHSKPRVQVGGTLGNTP<br>PANISYPLKVQGHTFNNVHVFRDFVSQGCASCGKIPTAYDQRNRHLTVYEGAKFTGSLDECEFICGECVIENK | Putative amidotransfer ase | Ceyssens et al., 2008 |
| gp25 | 74 | MTQVTLKRQVVIQMETDATRKYPFSRDTLDKIQSIRRVKEQELNDANPDEEFLVPAPVVIAEAIDRLFEDYFE | | |
| gp26 | 295 | MFYIYKGSRASSGAVNLRNALGARILRSEGSTYRGRAGTAVINWGTVGAEARRLQGIAPVFLNDPAMVARCT<br>NKLDFFRHFEANAPHLIPRWTERWEDALGVLNISGRMYARTDLNGHSGRGIHLVCTVNDAEVAAIDALRRQG<br>HYPVHIYGHTHIPDVITRAQLFTQGIVGKRTEFRAHVIRGEVALLQVKLRRTANDTVTNEGQSIVRNVAGGWV<br>YGVNDAMGRDGAEQAMSAAAEAIQVAGLDFGAVDIIYQHATSRAFVLEINTAPGLDAEGSALEAYVKGFNKI<br>FEETI | | |
| gp27 | 133 | MAVRVFVYGTLLSGLYNHYLLEGAEFVGNAVSCERGLMYSAGGFPILSFASRADLIVGEIWQLPEGEAGDEML<br>ENLDALEGYPGWYDRTLKDFRINGERIKALVYHQDSHMAMDIVKDGDWKAHLAKRQGAV | | |
| gp28 | 126 | MNEMTVDKAVEVYRDTPNTFGHQELHAQKMLLKEILGLVASQRHLQDSIEVSKIPEASDSPETSYGGYCDESI<br>GIRFMWERLKKIEDRLRELEEVYGTFVTTPYKTLPGNVNAVPSLVLKSQLEG | | |

| Name | Length | Sequence | Function | Source |
|---|---|---|---|---|
| gp29 | 570 | MKKIIGDTACPGCRAKGGDKTGNHLILFVDTEKGTRFGSCNRCGHYEVLEEGFKVPERREKSEEDIIHEVNEVLEYPIKALDTRKISKSIAERYGVRVGLSQENGEDVIEHYYPRTRDGEYRAFNVRILEPKAFYYRGSPKGGVDPFGYNTLRHKDMGHLRLVICEDELSAMSVAQIMESKLPEKWKHLRQASISWSSGVGSAGRDIAFLKESGVLERFNEVIYCHDADDEGRKSVEKVRALYPECKFVELPLKDANDMLMRNRGDEVYQMIRFGSKVKSPDCSVTVDEVYAEALEPPKWGKSYPWEGLTNLTYGQRDGEIIGVGGGTGIGKTLLAHEIAAWNCIEHGENVGTFLLEEQVAMTLKNIAGKVANVPFHRPDIEWDEQAFKDAAGKLRGKLFMWKNKGQNDWDHIKECIRFWAVAMDVKTILLDNMTAMTNHLSPSEMNTEIARICTELAGMADELGLRIFIFSHLNPPKGNRTHEEGAEVKESQFTGSRAMQRWCQLMIGFERNKQADGEEKHESRIRVIKDRNYGNTGLVFTKYNPETGRLVEREGSYDEVPADDDTPI | Putative DNA primase/helicase | Ceyssens et al., 2008 |
| gp30 | 170 | MTPQFDYVIYDLEGDGLFNEVTRLWCAVAVDIPTGVVRGFRPEEMDVFYRIIAHAKFVVGHNILDYDNRVLEKLHGIIIPPDRSYDTLVASRLTWPDRPQGHSLGAWGRFLKCHKGDFNDFSKFSEEMFEYCLQDGVVSHALFNYLLRVLGMTWQELVEWRTVDWLKSE | Putative ribonuclease | This study; predicted function based on homology |
| gp31 | 99 | MAKKRVRNYKRERELAIRRGETGVGSKSGDAQRHRARRKVEKRLGRKLGADEVVDHIKRVKDGGGNGDSNLRVRSHSSNAADGGRVGNRKAKGIRKKK | | |
| gp32 | 78 | MFNRKLSISNILNSFDKVLVNLKTFIQESSEESERIYNEISLLKAERTQVMQDNLKAQKVLANLEELLGGKSEEVSS | | |
| gp33 | 64 | MKKYRVNVGFQDTKVFNADFYRIELDIIRFFAGDSDANPMTVRANEIGAVRGWVSVEEINDGE | | |
| gp34 | 54 | MASKKESLEDQARKEIALEKEFSGSWGGPEIDADDFPLGSACGLDPEVCESCQ | | |
| gp35 | 95 | MPIALQVLVIFWSAFFQVFLLGLNSKLLRDDKIKAGFVVSWCITLAQFAYIKAVGSSHLDIGWFIFVSGWGGALGITSAQYFYKWYDRVFHKKA | Putative holin | Ceyssens et al., 2008 |
| gp36 | 170 | MSDHVSYSKHVRGKYLCNMASALHKSMEVQRTNIRKFLSSPHITLREKRRVFLSLPEGFLGVSYFTGSHLNLSSYSDRRNTRIRDKDMSLYDDFYVDRGTQLDPRDALFANKEAKEYEFQFLRKKWGTDFELSGEDLENVKGCQRMIDLSWYLVDLACERGCSYFVFDW | | |
| gp37 | 66 | MSKIKSILMERVDDFLLKQVAVAFLEQQWKLDRSGTVDYLSYLGGVSHEAVETVVENLAERLKGD | | |
| gp38 | 546 | MDWRKSLFVEHKVADIISRQSKRGVYFQTQRAKWLIHVLSERILKIDLEAVPQMPMMIVRAGAFSKPFLKSGKPNQRLQSLWQRLGHFEVSGPFSAIEYVPFDLGKTAKFKDWMLDQGWIPDQWNIKDITVGTDGKKLRGSDLNESLNKYIEDLRQSKSGRLRMKLQGIIPGKTTIGEVKRKLEKQRKVLTTPKMTEESMDTVQGDLGKLVMQRMVWAHRRSLLQGLVDQVRPDGRLEGSANPCATPTGRMRHRVVVNIPAARSPFGPEIRGLFQGTPEAGEWKWTVLRRDLGENERVRPFTNIVEELKKGKWKTVGKYRVYVPANQLVFVGYDGAGLELRMLASYINNPEYTKEVVEGDVHTANQIAAGLPTRDDAKTFIYAFIYGAGDAKIGTIIGGTRADGAKLRAQFLEANPDLAALIERVKQEAERGYLEGLDGRKLTMRRSESGDVMIHKALNTLLQAAGAIVMKWAMVLLDERVRRLNLRAWKVLDIHDEGQWECHPEDLKALREQMEVCVRDAGEILGVNCPLASDSIAGRSWKDTH | Putative DNA polymerase | Ceyssens et al., 2008 |

| Name | Length | Sequence | Function | Source |
|---|---|---|---|---|
| gp39 | 187 | MSKKVSQRFTFPVAKLIFPYIVTPDTEYGEVYQVTICIPTKEEADKLVQQMESKDVRLKDTIKYQERDGEYLFKVKQKKHVDWMQDGERKSAVMKPVVLTSDNKPYDGPNPWGGSTGEVGILIETQKGPRGKGTMTALRLRGVRLHEIVSGGDGEDDPLFGGGFAEEEDKSEDVFDEVFDDEDAPI | | |
| gp40 | 168 | MTKTLLSKGLADHESGVCRSGCPYCLIEFEKVWGVRVVSSTAASNNKVEVDPNGIKPGEPGAKLDSGKVDVGIIFEAFPRALYAVAQVANFGASKYSRGGWRHVENGIQRYDAAFGRHLLERHKGKTLDPQSKLPHRYHEVWNALASLELVIQQEEESNGTSVGPKG | | |
| gp41 | 95 | MELLLDPRAKTVPSNYSVMGIDVDLGLPPGYSLTEEAMDKAKRQESEYYDWKGYEALANPVIEHPEYRAKGEAFALRVFWEEKLKESQVVEEVS | | |
| gp42 | 43 | MIEERFWSKVDKTEGCWNWTLGFPYNAVQNIRYGRAWAQCWE | | |
| gp43 | 277 | MSKEHIFDIQVEKPWPEEEVHKLVDDKVEQIIKRVNADECEIYLTGQGNFRLELAKIKQYKGTRIGLEKPHHWETVSARLKDKWGAITFHGIEADDWLGIRGTEEGDNFTACSRDKDIRQVPGCYHYSWPCGDSQPELGPFQVDGLGTVSASWRMYGVKKPQKSWKLEGNGTAFLYGQLLVGDSVDNIPGLPGTGPKTAADLLGELSNERDLFAACAYAYQQKYGDNWKEYLVENFRLLYLIRDRSWLDIQQSGNEYHCSLKKHWEIPYDDEDLFY | Putative 5'-3' exonuclease | Ceyssens et al., 2008 |
| gp44 | 328 | MTTKIYSIDEALSICEGLFKILETTGFTDYKVAGGFLRDADNGVAPKDIDLYVRRPRVEAIGRRVSRVFGPQMVPCSDDELEVEVIRFYNKLGHKKVRTGSGELRNGYPEGFSVWESIGVDLPVNLVVSTDSHPAEFDMGICEISCWPIGTLRRNRSGEIYRSRAYEWDKENKCITINRMVDPLLESGNEITVGQIERLVNHVSRVKAKYPDHRVCIGDWIWMLLRSRSVYAGSAMDFIVRLQEEGLIGKAGEILQNQTEVIDWDEVRQRNREDRPRDDALDAVQAVPGTIRHRAQVQAGLQGIDITTLWIDEESVGRGQGVLPRGV | | |
| gp45 | 80 | MVEVKGYFQEASEASKYIWVREALPPDTELVFIFERPNTACHWLSKRKDGTKQSMAEWAERNGFRWFTLETFKESFPNE | | |
| gp46 | 254 | MSKKYNEDTLVIADTQVRSEVNIDHIGNLGEWIARNRPKRIVHIGDHWDMPSLSSYDRGTAKIEGRRVLADIQAGNDAMRVLLDPLRRLQQHQASCKKRVYRPEMHFFIGNHEERIKRYENSNPALQGFIGYDHFDLSDWIVHDFLDVGVIEGVAFAHYFYNPNSGRPYGGSAEHRLNKIKRSFVQGHEQGFKYHIEAVGKKRIHGLVVGSFYTHDESYKGPQGNDHWRGVALLRNHKDGEYDLKLMSVEEFL | | |
| gp47 | 73 | MSKFLPDLYYIKSEHDFGQRGLAFKTPISAELWLDMKFGKGGAEEGLKRGMYSIEVLEILYIPSVHLPDILG | | |
| gp48 | 66 | MGRKLEVGDSVVFLVHRNTSSHLAIGTVDGFTPKMIRIKCPTMSWTIDSEYVLRSSDKVVYYDKG | | |
| gp49 | 69 | MTKAELEKALEETQSALAKAEAKAFSFEELAEEAKRQIEFLEGMLDLVDLRASVFYGDWRGYAERSKG | | |
| gp49.1 | 32 | MQKDQKGKGGFPWTYIAVAALFALLVYVGYS | | |
| gp49.2 | 24 | MLLLTLGEISRLLIEVLSWAGSL | | |
| gp50 | 63 | MKYRVQQLGKRLFYIQKRFLFMWRTVTDSGGCFMAFYSLEDAEEFIRDNLKYRDSPVYHPVE | | |
| gp51 | 119 | MAGYIADNDGNSVFVIQSKAIAVRNEDGTLRQVIGPKDFEPVKWEDIEGKPKLLEVGAKASEAKPGNWKPKASEVSGLYEAIAASIQDKISEIPVASYTDTHEALVDKFNTLLNALRS | | |

| Name | Length | Sequence | Function | Source |
|---|---|---|---|---|
| gp52 | 296 | MATYVETDAAGPGGRAYVVDVPKVFRRNADGDLIEITSDAGRVNTNIVTATEFLPDYYAVKENAALVVENVT<br>ASFPENPSEAVIPSVLVNTLVGAPGETSGGNRDPGHVIGLFSETMIEAPCGTAFGSEFRVDPRREHLDVYVAIK<br>HVIGPDDQNGGTIGDYIIEQFDDMRGPVQNIGSLQQNYLDPRLVTTHLGGNVVNTQQLTQNITLTKQQSGW<br>FFMSQGTTEITVTLGPDVTPGCHFHFIQGSAAKIKFAVSPDKAWYAKGNQTETDGQFGECTIRVYPFGGTVGT<br>FKSAA | Decoration protein/head spikes | This study |
| gp53 | 1056 | MALSPELKAAIDAELAEVKGLDVAAELASLNEPEVIEEAPQEVAPATPEPAPDLSALVTPADPNSISSAIGRGVD<br>TMQSNIGGTIATLGELTGSDYLKDYGTQMAEENAQEASQYGTPDVRSFADIQDIPSIGSFLKNNIVEALPSMAP<br>VLAGGAAGAKAGSVFGAPGRMGGALIGSFLSSMGINVGALSNQMKELDPDQSNPWTAVIGGAGLSALDTA<br>GAGVIAGPLLKHIGKDGAYQMLVQSGLPKQTAIEAVTQAGKHAAVSGVAEGVTSGAQQALQDTIAYDAVGQ<br>VQTPEQFMDNLLTAAFTGTAIGTAGGAVSSGLDTLGRHQDSAGSAVVDPNAPEAPPKASEFEPRGTVKKAW<br>DALGNEATSLLEPLAKASPIAREFSETFRADMSGKRASGKTIFEDQELQAGKWNSELDNIFEGKSSKEIDRIISDT<br>SAGVNTPEATRLRALMDDVRNEAVNRGGMSVGTIPNYMPFGLSPEKVQTPEFLNDITPYFQNRQAAEDAVA<br>NWLAEVSDDTRGNTAPEVNRLVTQNQQTGAWEVDPRYRIQGDPDTLRGRFAQSDAVPKYGQLEESRAFGS<br>VPQEILNKYSLNDTPKKRLQEIRDYFEGASHRIAFTERFGINGEKANAKIASAVAEAQRAGKRVTKEEVDRMYD<br>LVDAYNGMHGRIKDPNLKKLAAVTSGALVLSRLPLAGFSTLTEFSLPFAKAGVMPTLGAVLPTMGEVVRQAAR<br>RIYSGVPKSETGRFMSDMNHTLSSATSLMADRVGAEVFNTTVQKAIRGQFLINGLSILTHVNRVFATETAKRVY<br>QNNLMDLAAGLPFSSANGALKVAQLREMGVNIGIQQDALKLVSPATPSEVLMANNVKTLAMRRFVDQVVL<br>DPTFADKPMWMSNGNVQMFSLLKGYPAAYGNIILPMFRRRMSPHFAGSWTNAGMGAAGVAFTLGLMM<br>SLGYLQDELRQLAKFGGSSREDTRSPEQRMMDVVMQQMPLQASMIYDMLTGYRRGTTPAEVLLGPVAGAA<br>TEGAMAVGKTIASFGDDPSAGEIWKFLYKQTPARPFVAGMEAMEDALDL | Putative inner membrane pore forming protein | This study |
| gp54 | 506 | MASMAYEGSPIRPSILRAAQNELDMARIARNKLPLAVEGSGLPDRVRRAAQAALENPDRWSRAAQEVAPAA<br>EATGRGALGRIAGILGGPVSVGIQAAVTPGELGDAERTRAEEMAQASQAVENMGPEVAQEANQWAQGVG<br>QRAAQNATGGPTGAELLSYGVTPNQPSIEPEITPEVASEAGAAVADEEEANRQVIQQGAAEGLRTGAVSRPE<br>MAQAVVEADAQREGVELKPQELKNRVNEELTQMRTMDNDDLSRYVSYALIGTGLLASAIDKTGKAGDMFA<br>ASYERQLDRNLQAGINQQKMAAAAADRQIKEKDLERKVAKDAADVRLGEGNLEVKKGTLEETSRKNTGLLDR<br>WAEEAARGRANLALTQRGQDMANQRAQLQAETTRRGQDMSQENAQLSSAVRLKTAKISAQARQAAAKAA<br>RGEPVTTKDALGILSEVSGSQALGGKKLGKTAQQAIAQTLRNEMRANPGANPIGIIQREAAKLQPTGNWFFG<br>GDLDYPAPTR | Ejection periplasmic tunnel | This study |
| gp55 | 127 | MASLPQKLFAIGQNIGGGQERVQLSRQGSYRPTHLGTMQSGESSGQSNPFGAMGGAALAALLGQGSEPSSE<br>AVPSFSVEGARGASERGAAEVAAGMGAGMGILPSAEELGFGQQPKSGILSKLFGG | | |
| gp56 | 314 | MNFLQFNVKRLMGFDVEDALDSRAYKGGSKKQKTTSVSTPYQQGNYNELLSGASDWLHNGGFDPNYGGDP<br>NFDPVADQNALQLGGIEGLGGLGGALQSLLGSSGVSSLADYLGPYDPNKTGLANAIGAANEQMQWDFDTTV<br>RPDLRAGATNAGQYGSSRAGVAEGIATARLLQNQQNNASQLAFQDQQAYNQNRLNTLGNLSAIAKGLGSG<br>NAMQVDAGSMLQNQEQQEINGALQKWAYENNVSLNDLLAYKELISGDMGGTNVTTSSGGGGGGGLGSAL<br>GALGGASLGALFGGPAGASVGMNTGGRVGGLLF | Putative outer membrane pore-forming protein | This study |

| Name | Length | Sequence | Function | Source |
|---|---|---|---|---|
| gp57 | 145 | MIHLITRENIDLLPTVVPALARAFNRTDLGKFWDFEHLVHSLVNYEAYVFYQEESGYAGVIQVSQAPLGNILHFF<br>WSGKMPGNETPVDYSEVDDFLGQFAQRVNCRFIQCEGRRGWKPTLEKLGYTEDSVSFYREVTPDELPPI | | |
| gp58 | 230 | MSQKYSPSIPPQEEEELLPFLNEEFVRVGQTLNDLADGYWGVSMEPPKKLKPGTVKYFAPGVVGPVSGIYHYD<br>LDNQWRLAGTKPKDLPGDFILFTPQNNHQPMGTCAYRMNTAKDEVWITMLMSGGNYTNGATVLDLPQAY<br>WPPAELFIPAYSSIIPAQSTITYPPPSDPNAPPLDQVFDVLNRATIQTGVVNQAMFKITANGRVLIQGIPQGAVF<br>GGTFTFPLVVTP | Tail needle | This study |
| gp59 | 514 | MALERQEVKNPTGIVTDIAPADLPLEKWSFGNNVRFKNGKAQKALGHTPIFDTAQAPILDMFPFIRNNIPYWL<br>LCGEQRMYLADGTTVVDVSPGGHSASVTSRWSSGSFNGVIFANNPSNYPYVLMPQNSGFIPMPNWPANTF<br>AKRMKSFKNFMIALNVTQNSVEMPQMVWWSTSADAGGIPVSWDPTDPTKDAGQNTLADTNGAIVDGVK<br>LRDSFIIYKEDSVYSMRYIGGLFIFQFQQLFNDVGILGPNCAIEFDGNHFVVGHGDVYVHNGVQKQSVIDAQV<br>RKFFFSDINPDNYQRTFVIADHVNTEMWVCYSSTRSEPGKHCDRAIIWNWKENTWSIRDLPNVLSGAYGIIDP<br>KVSNLWDDDPNPWDTYTSVWGEGSYNPAKSSMIFSSFQDKKLFLFGNNSTFSGQNFVSTLERSDIYLGDDR<br>MMKTVSAIIPHITGNGTCNIWVGNAQVQGSGIRWKGPYPYRIGQDYKIDTKHVGRYIALKFDFSSEGDWYFN<br>GYTIEMAPKAGMR | Tail hub | This study |
| gp60 | 216 | MGLEVATYINQLVPTNPTGSDLKSFGDDHLRLIKSAIKNTFPNISQAVTVTAAQLNAVADTTQYVKPGMVIM<br>WAGSLAQIPAGWKLCNGVGTTSNGIPVPNLIGAFPWGIDGSSQAVGTRGGSANIVWDGFTEGTALTLAQIP<br>AHTHTWRSRGATTLTGSAGDSGALTGGSGNAANTNLETGPAGQGQTHNHAVKINMPLGNIPPFCSVFFIIKN | Tail fiber protein | This study |
| gp61 | 83 | MQLLTDNLGSIVSALVVSSVVTVGSSVVGQNVHQALLERNIQATEKLTKAVSDLQISMAIFGERYVTRKEMKE<br>EIKEAKNGS | | |
| gp62 | 64 | MPIFLKTLAINLLATLFPTKVIAKSIVTGAKALAEKTNSKVDDEFVKVLEDNLKEKTEDATSN | | |
| gp63 | 209 | MATINNVTDLAIAAIQWSDRQDLTQELLMLFIGNTTDRLNRLLRVRENEHFETLMAFGGGIEIPEHFVALRSIT<br>GDSLIGGRTLQYITQDIFTHYVNYNYQPQGVTYYTRLGNFWRVFPVVPDGAPFIVNYWTVLPELSLANPTTWA<br>LTKYPQIYLYGVLEQIYLYTMDEARSQFWGQKLERAVMELQNEENAADFASTRLAIKDIER | Head-to-tail adaptor | This study |
| gp64 | 107 | MSIQSTYAIDEDLGLFIRKDTQDVESILSANAEERHSGVNSNRKDNMRKVASIPLVVVEALRNRPMAEGGPIDL<br>NLIGCDPDHAARFTRWLNDRDNYRMRTSEARV | | |
| gp65 | 318 | MATPTNAVSTVEINGKREDLIDIIYNIAPYDTPFMTAIGKGVATAITHEWQTDELRQPGKNTRVEGEDATIKAG<br>SFTTMLNNFCQISDETLQVTGTADKVKKAGRKNELAYQLAKKSKELKLDMEYAMVGAPQAKIQRNTTTPGQ<br>MANIFAYYKTNGSVGANGTLPTGDGSDTGTAGDLRLLTEDMLLNASEAIWRNGGQANSIQTSSSIKKAISKN<br>MKGRATEITLDASDNRIAQTVDVYESDFGKYTIRANRWFHENTMFIFDPKMHALCYLRPFFQHELAKTGDSEK<br>RQLLVEYTLRVNNEKSGALIRDVVAQV | Major capsid protein | This study |

183

| Name | Length | Sequence | Function | Source |
|---|---|---|---|---|
| gp66 | 331 | MDLENQGMDEFEAAEAFGDLLGDDLLAPGDETVDGKVVGEEEEEEVVDEGTPEDTEEVESEEEEEGSEEDPE<br>VESEEDPEDKLFEIPIGDEVYEVNFEELKSGYLRNEEFVTRQAELEDQYAAKFEELDAERSNLLAELEQYAVTAIA<br>GANQYDNVNWEQLKAQDPEKYQNLRLEALEARDRAQALIKRRNDIKAMQDKRAEIVHSAYVKRQTELAKKLI<br>PEMTTDESWGDKIVSYGKSIGYSEDEIRGISDARQLAVLDAARKWAESQVRRKAALEKKEETELPAAVKPAAR<br>RAEASEGSKRVKAARANLRKDQSVEAAAALFSSLDIF | Putative scaffolding protein | This study |
| gp67 | 81 | MGERFGYSLVLTDAAQQMVRENVLPELFRIVQEEIEGEWKATAPKDSETREAIYHELHALNRVQLRIQAILDSI<br>TLKEYK | | |
| gp68 | 707 | MAKRRRKIKPMDDEQVLRHLDQLVNDALDFNSSELSKQRSEALKYYFGEPFGNERPGKSAIVSRDVQETVDW<br>IMPSLMKVFTSGGQVVKYEPQTAEDVEQAEQETEYVNYLFMRKNEGFKVMFDWFQDTLMMKTGVVKVYV<br>EEVLNPTFERFSGLSEEMVADILADPDTEILAQSVDEDGTYSIKIRKDKKKREIKVTCIKPENFLVDRLATCIDDAR<br>FLCHREKYTVSDLRLLGVPEDVLDELPYDEYEFSDSQPERLVRDNFDMTGQLQYNSGDDAEANREVWASECY<br>TLLDVDGDGISELRRILYVGDYIISNEPWDCRPFADLNAYRIAHKFHGMSVYDKIRDIQEIRSVLMRNIMDNIYR<br>TNQGRSVVLDGQVNLEDLLTNEAAGIVRVKAMNSIMPLETPQLSGEVYGMLDRLEADRGKRTGITDRTRGLD<br>QNTLHSNQAAMSVNQLMTAAEQQIDLIARMFAETGVKRLFQLLHDHAIKYQNQEEVFQLRGKWVAINPAN<br>WRERSDLTVTVGIGNMNKDQQMLHLMRIWEMAQAVVGGGGLGVLVSEQNLYNILKEVTENAGYKDPDRF<br>WTNPDSPEAQQAKAIREQKEAQPKPEDIKAQADAQRAQSDALAKQAEAQMKQVEAQIRLAEIELKKQEAVL<br>QQREMALKEAELQLERDRFTWERARNEAEYHLEATQARAAYIGDGKVPETKKPSKAVRK | Portal protein | This study |
| gp69 | 483 | MDTQERLRNLVKELAERQKYFRMNQYTPYGWQEKFIAASSNCAQLLAMTGNRCGKTYTGAFIMACHLTGRY<br>PEWWTGRKYDRPVNCWAAGISTDTTRDILQSELLGDWKNPEAFGTGMIPKEDIVETIRREGKPGCVQAVVV<br>KHASGGLSSLIFKSYEMSQDKFMGTAIDVIWLDEECPKDIYTQCVTRTATTGGIVYLTFTPEHGLTEIVKDFLQD<br>LKPGQFLVHASWEDAPHLSPEVKEQLLSVYSPAERRMRAEGVPMLGSGVVFPILEEKFVCEPFQIPDHFHRIIGI<br>DLGFDHPNAIACVAWDPEKDKYYLYDERSESGETLGMHADAIYLKGGHQIPVVVPHDAFKHDGATSGRRFVD<br>LLKDDHNLNVVYEPFSNPPGPDGKHGGNSVEFGVNWMLTRMENGDLKVFNTCTNFLKEMKMYHRKDGKII<br>DRNDDMISATRYALLMASRHARPGAVRNSGYYRSDTAKLTPDWFGSIV | Long terminase subunit | Shen et al., 2012 |
| gp70 | 114 | MQLPGEDGPITLARAETWLEQDSKAAYDAAQGQVSQLPFCTPELFDALVSVNFQLGTAWTKKFPKTWNLLK<br>SGRFDEAAWEAEDSLWAKQTPVRVRDLQRALWRASTVGDKVV | Putative lysin | This study; predicted function based on homology |
| gp71 | 153 | MSDEKVVSIGAAPLSAKEKLDLYCEALADGMNKTQAYIAAGFSPNHAQRNVAAYHRKHAEYINAFISERIGSH<br>VPMALRVVVEIASNPEEKGGIRLKAAQDILDRGGFGAKQKLELTTKNVEDMSAEDLISEVRRILDEEPELAKVIS<br>FPAS | Small terminase subunit | Shen et al., 2012 |
